# The *Toxoplasma gondii* effector GRA83 modulates the host’s innate immune response to regulate parasite infection

**DOI:** 10.1101/2023.05.31.543158

**Authors:** Amara C. Thind, Caroline M. Mota, Ana Paula N. Gonçalves, Jihui Sha, James A. Wohlschlegel, Tiago W.P. Mineo, Peter J. Bradley

## Abstract

*Toxoplasma gondii*’s propensity to infect its host and cause disease is highly dependent on its ability to modulate host cell functions. One of the strategies the parasite uses to accomplish this is via the export of effector proteins from the secretory dense granules. Dense granule (GRA) proteins are known to play roles in nutrient acquisition, host cell cycle manipulation, and immune regulation. Here, we characterize a novel dense granule protein named GRA83, which localizes to the parasitophorous vacuole in tachyzoites and bradyzoites. Disruption of *GRA83* results in increased virulence, weight loss, and parasitemia during the acute infection, as well as a marked increase in the cyst burden during the chronic infection. This increased parasitemia was associated with an accumulation of inflammatory infiltrates in tissues in both the acute and chronic infection. Murine macrophages infected with Δ*gra83* tachyzoites produced less interleukin-12 (IL-12) *in vitro*, which was confirmed with reduced IL-12 and interferon gamma (IFN-γ) *in vivo*. This dysregulation of cytokines correlates with reduced nuclear translocation of the p65 subunit of the NF-κB complex. While GRA15 similarly regulates NF-κB, infection with Δ*gra83/*Δ*gra15* parasites did not further reduce p65 translocation to the host cell nucleus, suggesting these GRAs function in converging pathways. We also used proximity labelling experiments to reveal candidate GRA83 interacting *T. gondii* derived partners. Taken together, this work reveals a novel effector that stimulates the innate immune response, enabling the host to limit parasite burden.

**Importance:** *Toxoplasma gondii* poses a significant public health concern as it is recognized as one of the leading foodborne pathogens in the United States. Infection with the parasite can cause congenital defects in neonates, life-threatening complications in immunosuppressed patients, and ocular disease. Specialized secretory organelles, including the dense granules, play an important role in the parasite’s ability to efficiently invade and regulate components of the host’s infection response machinery to limit parasite clearance and establish an acute infection. *Toxoplasma’*s ability to avoid early clearance, while also successfully infecting the host long enough to establish a persistent chronic infection, is crucial in allowing for its transmission to a new host. While multiple GRAs directly modulate host signaling pathways, they do so in various ways highlighting the parasite’s diverse arsenal of effectors that govern infection. Understanding how parasite-derived effectors harness host functions to evade defenses yet ensure a robust infection are important for understanding the complexity of the pathogen’s tightly regulated infection. In this study, we characterize a novel secreted protein named GRA83 that stimulates the host cell’s response to limit infection.

## Introduction

*Toxoplasma gondii* is an obligatory intracellular parasite of the phylum Apicomplexa that infects a wide variety of avian and mammalian organisms, including humans (1). This widespread parasite is estimated to infect one quarter of the U.S. population and as many as two billion individuals worldwide (2, 3). Although infection in immunocompetent individuals is typically asymptomatic, infection with the parasite can cause congenital defects in neonates, life-threatening complications in immunosuppressed patients, and ocular disease (4–7). After ingesting *T. gondii*, the acute infection is established by rapidly replicating tachyzoites that disseminate from the intestines to peripheral organs, including the central nervous system. This form of the parasite is ultimately controlled by the host’s immune system, but some of the parasites are able to switch to bradyzoites and form latent tissue cysts (8). These cysts persist for the life of the host and can then reactivate the infection during immunosuppressive conditions (4).

To control the infection by *T. gondii*, the host unleashes a wide array of innate and adaptive immune responses (9). These responses involve distinct cellular and humoral mechanisms aimed at dampening the parasite’s rapid replication, thereby avoiding infection-induced pathogenesis. The paramount immune factor against *T. gondii* is interferon gamma (IFN-γ), which orchestrates effector mechanisms that will successfully kill the parasite during the acute phase of the infection, as well as keep the latent stages quiescent (9–11).

*T. gondii’s* initial interactions with innate immune cells play a crucial role in determining the outcome of infection in the murine model (9, 13). The recognition of profilin, a protein associated with the parasites’ actin polymerization, by Toll-like receptors TLR11 and TLR12 located inside the endolysosomes, leads to macrophage production of interleukin 12 (IL-12) (14–16). IL-12 production is mediated by NF-κB, which is the key inducer of IFN-γ by natural killer (NK), TCD4^+^ and TCD8^+^ cells (17–19). IFN-γ is readily consumed and activates residing macrophages that are responsible for parasite killing through nitric oxide production (20), tryptophan metabolism (21), and GTPase disruption of the parasitophorous vacuole (PV) (22), among other mechanisms.

*T. gondii*’s ability to successfully invade and replicate within virtually any nucleated cell is dependent on a sequential release of parasite effectors from specialized secretory organelles called the rhoptries and dense granules (23). *T. gondii* tachyzoites actively invade the host cell and replicate within a host-membrane derived PV, which is able to escape host clearance, enabling intracellular survival. Upon host cell invasion, *T. gondii* secretes an array of dense granule (GRA) proteins into the PV. Some of these GRA proteins regulate vacuolar functions, such as maturation of the PV and nutrient acquisition (24). Other GRAs traverse the vacuolar membrane to reach the host cytoplasm where they actively modulate host cell functions to optimize the parasite’s replication and succeed in colonizing host tissue (23).

Numerous GRAs have been studied in the context of the acute infection that contribute to the intricacy of the parasite and host interplay. Some of these GRAs restrict parasite replication and dissemination to limit the infection, while others allow for its persistence in different host tissues. One of these is GRA16, which binds two host enzymes, the deubiquitinase HUASP and PP2A phosphatase, resulting in accumulation of host c-Myc, regulation of p53, and the cell cycle which ultimately results in attenuated virulence (25, 26). Another of these effectors is GRA28, which is responsible for facilitating parasite dissemination during pregnancy by inducing CCL22, a chemokine critical for immune tolerance during pregnancy in placental and monocyte-like cells (27). GRA28 is also the mediator of upregulated CCR7 and responsible for the hypermigratory phenotype exhibited by *T. gondii* infected macrophages (28). While these GRAs serve to enable infection, GRA24 dampens parasite burden by interacting with host p38α MAPK, leading to its activation and downstream stimulation of inflammatory cytokines (29). Similarly, GRA15 interacts with host tumor necrosis factor (TNF) receptor-associated factors (TRAFs) and activates the nuclear factor-κB (NF-κB) pathway to drive transcription of IL-12 (30–32). These are just some of the examples of GRAs that highlight the parasite’s diverse arsenal of effectors that modulate infection. Elucidating the full complement of the parasite-derived effectors that hijack host functions to favor intracellular survival, as well as those that regulate host defense, is important for understanding the intricacy of how the infection is regulated. A mechanistic understanding of the processes is also important for the establishment of efficient prophylactic and therapeutic protocols to effectively treat *T. gondii* infections.

Herein, we characterize a new GRA protein named GRA83 that is secreted into the vacuole in both tachyzoites and bradyzoites. Disruption of *GRA83* surprisingly increases parasite burden and overall susceptibility to *T. gondii* infection in mice. This increased virulence correlates with dysregulation of cytokines in acutely infected mice, indicating that GRA83 modulates innate immune mechanisms. This dysregulation of cytokines is linked to a decrease in NF-κB activation in Δ*gra83* parasite infections. Removal of GRA83 also results in a substantial increase in cyst burden observed in chronically infected mice. We also report potential *T. gondii*-derived GRA83 interacting partners through proximity labeling using GRA83 as bait. Together, we determined that GRA83 enhances the host response to *Toxoplasma* infection and therefore plays an important role in the interplay between the parasite and its host.

## Results

### GRA83 is a dense granule protein that is secreted into the PV

TgME49_297900 was initially identified in our MAG1-BioID studies, which involved using *in vivo* biotinylation of the bradyzoite PV to discover novel dense granule proteins (33). TgME49_297900 was also predicted to be a GRA protein by the recent *T. gondii* hyperLOPIT (Localization of Organelle Proteins by Isotope Tagging) analysis (34). OrthoMCL predicts that TgME49_297900 is restricted to *Toxoplasma* and its closest relatives (*Neospora, Hammondia, and Besnoitia*), but is not present in other coccidians or apicomplexans (35). TgME49_297900 encodes an ∼83-kDa protein that lacks a signal peptide but contains a predicted hydrophobic domain (residues 74-92) near the N-terminus that could play a role in secretion or membrane association (Fig. 1A) (36). Similar to other GRAs, TgME49_297900 contains intrinsically disordered regions and has no discernible protein motifs or domains that suggest its function (Fig. 1B). In agreement with this, AlphaFold provides only a very low confidence structural prediction for TgME49_297900 (Fig. S1) (37, 38).

**Figure 1.**
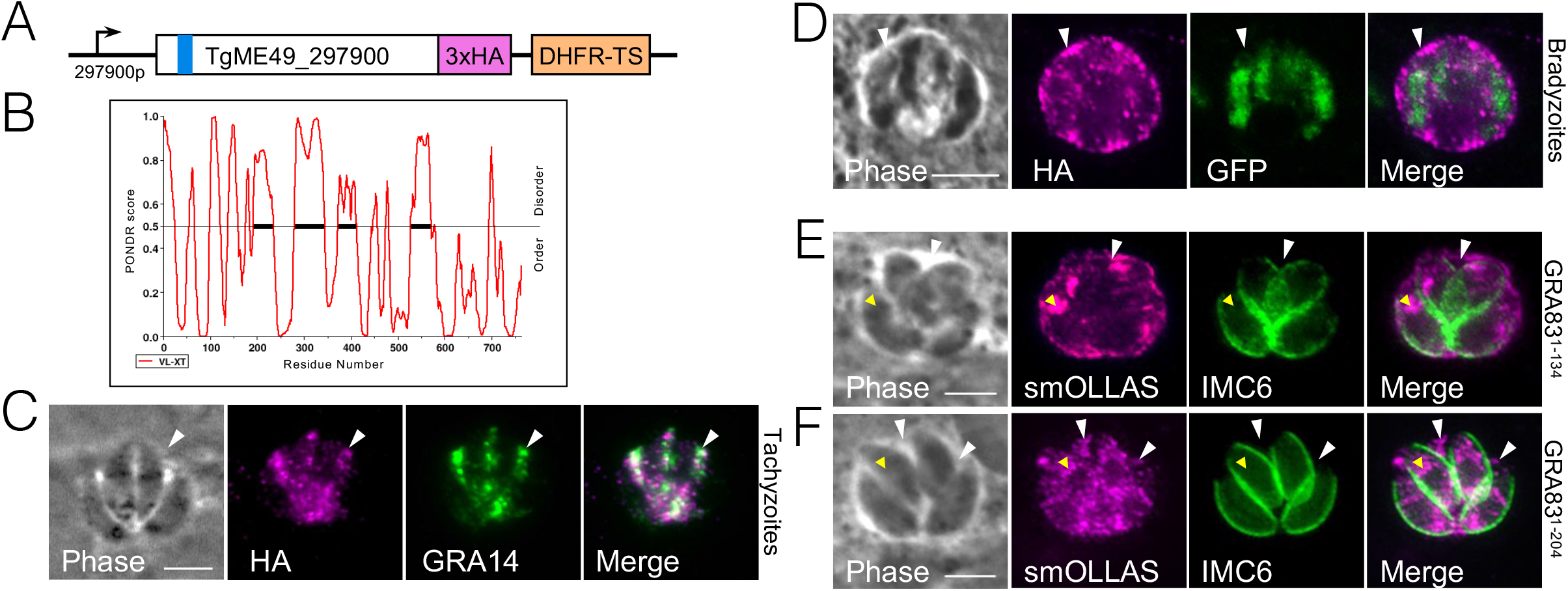
TGME49_297900 is a dense granule protein that localizes to the PV. A) Diagram of TgME49_297900 showing its N-terminal hydrophobic domain (blue bar), as well as the C-terminal endogenous 3xHA epitope tag and the *DHFR-TS* cassette used for epitope tagging. B) PONDR Protein Disorder Predictor predicts GRA83 is 40% overall disordered. Scores higher than 0.5 are considered disordered. C) IFA of GRA83^3xHA^ parasites demonstrating protein accumulation in the vacuole between the tachyzoites (arrowhead) and colocalization with known dense granule protein GRA14. Magenta: rabbit anti-HA; Green: mouse anti-GRA14. D) IFA of GRA83^3xHA^ *in vitro* switched bradyzoites shows robust staining of GRA83 in the bradyzoite cyst and periphery (arrowhead). The parasites express cytosolic GFP confirming bradyzoite differentiation. Images were taken after three days of growth in alkaline stressed conditions. Magenta: rabbit anti-HA. E, F) IFA of GRA83 endogenously smOLLAS-tagged after residues 134 or 204. The truncated protein is secreted into the PV (white arrowhead), while some protein is also present in the parasite (yellow arrowhead). Magenta: rat anti-OLLAS; Green: rabbit anti-IMC6. All scale bars are = 5 µm.

To confirm the localization of the protein, we used endogenous gene tagging to add a C-terminal 3xHA epitope tag to TGME49_297900 in Prugniaud Δ*ku8*0Δ*hxgprt* parasites, which also express GFP under control of the bradyzoite specific *LDH2* promoter (39, 40) (Fig.1A). Immunofluorescence assays (IFA) show that the tagged protein traffics to the PV lumen and colocalizes with GRA14 (41) in tachyzoites and thus we named it GRA83 (Fig. 1C). Because we found this GRA in our previous data set that is enriched for bradyzoite proteins and its expression is upregulated in bradyzoites, we differentiated the tagged parasites to bradyzoites *in vitro* (42). Successful transition from the tachyzoite to the bradyzoite stage was confirmed by the expression of GFP. IFA of the bradyzoites demonstrated that GRA83^3xHA^ localized to the PV and the periphery of the cyst (Fig. 1D). We also tested whether the N-terminal region containing the hydrophobic domain is sufficient for secretion into the vacuole by integrating a smOLLAS (spaghetti monster *Escherichia coli* OmpF linker and mouse langerin fusion sequence) tag after sequences encoding the first 134 or 204 amino acids of the protein (43). IFA of these parasites showed the N-terminal region can direct secretion into the PV (Fig. 1E and F), although some material also remained within the parasite, likely in the ER. This is most likely due to misfolding of the smOLLAS tag fusions which partially disrupting trafficking, but it is also possible that more C-terminal regions of the protein play a role in secretion. Together, this data demonstrates that TGME49_297900 is a GRA protein that is secreted into the PV in both tachyzoites and bradyzoites and that the N-terminal region containing the hydrophobic domain functions in secretion.

### GRA83 does not affect parasite fitness *in vitro*

To assess the function of GRA83, we disrupted its gene from the 3xHA tagged background using CRISPR/Cas9 to obtain Δ*gra83* parasites. The deletion was confirmed by lack of staining of the tagged protein by IFA and PCR verification (Fig. 2A and B). We then complemented the knockout with a full length, 3xHA-tagged GRA83 construct driven by its endogenous promoter, which was targeted to the uracil phosphoribosyl transferase (UPRT) locus (Fig. 2C) (44). The resulting complemented strain was named GRA83c. IFA and western blot analyses showed that complementation restored GRA83 localization to the PV and approximate expression levels of the protein (Fig. 2D and E). We then evaluated the growth of Δ*gra83* parasites by plaque assay and found no significant difference in plaque size between GRA83^3xHA^, Δ*gra83* and GRA83c strains, demonstrating that this protein does not contribute substantially to parasite growth *in vitro* in HFFs (Fig. 2F).

**Figure 2.**
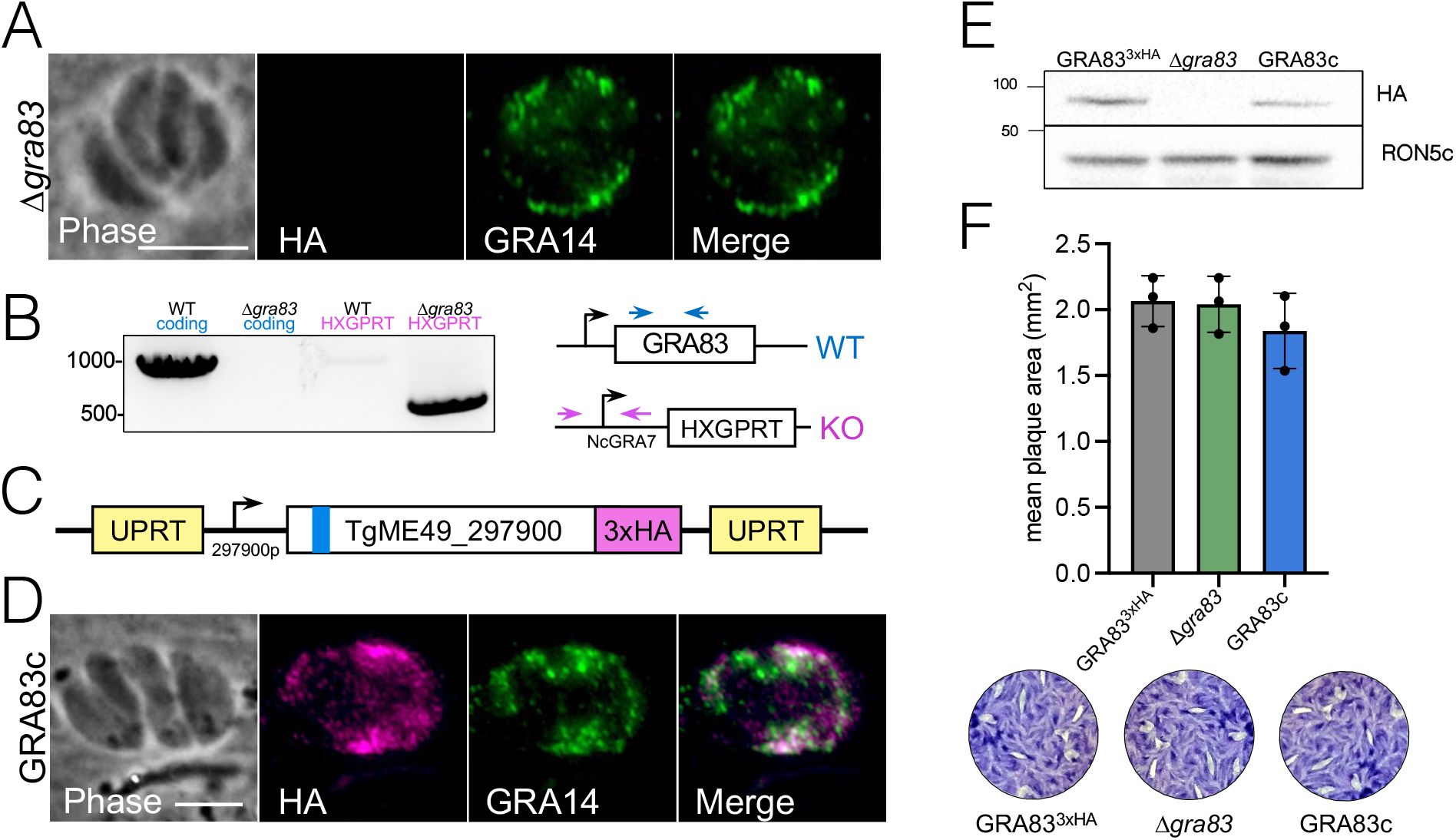
GRA83 is not required for growth in HFFs *in vitro*. A) IFA of Δ*gra83* parasites shows the absence of GRA83^3xHA^. Magenta: rabbit anti-HA; Green: mouse anti-GRA14. B) PCR verification showing that the Δ*gra83* strain contains the correct amplicon for the replacement of *GRA83* with the selectable marker *HXGPRT* and lacks the *GRA83*-coding amplicon. Diagram illustrates the positions of primers used to amplify the *GRA83* coding sequence (blue arrows) and the knockout construct in the *GRA83* locus (magenta arrows). C) Construct for GRA83 complementation showing the full length GRA83 tagged with 3xHA driven by its endogenous promoter and targeted to the *UPRT* locus. D) IFA of GRA83-complemented parasites (GRA83c) shows restoration of GRA83 localization in the PV. Magenta: rabbit anti-HA; Green: mouse anti-GRA14. E) Western blot of GRA83^3xHA^, Δ*gra83,* and GRA83c parasites shows the absence of endogenous GRA83 in the knockout strain and approximate levels restored in the GRA83c strain. GRA83 was detected with mouse anti-HA; RON5c was used as a loading control and detected with rabbit anti-RON5c. F) Quantification of plaque assays depicts no *in vitro* growth defect of Δ*gra83* parasites cultured in HFFs. The mean of each triplicate experiment per strain is shown. No significance detected using one-way ANOVA test. Representative images of the stained plaques are shown below the quantification. Scale bars = 5 µm.

### GRA83 potentiates mouse survival and parasite restriction *in vivo*

To determine whether GRA83 is relevant in the context of an *in vivo* infection, we assessed the virulence of the GRA83^3xHA^, Δ*gra83*, and GRA83c parasites in a C57BL/6 mouse survival assay. Surprisingly, we found that the Δ*gra83* strain was significantly more virulent, as most of the mice from this group succumbed to the infection at 10-12 days post infection (dpi), while all of the mice infected with the GRA83^3xHA^ or GRA83c lines survived the experimental period (Fig. 3A). Accordingly, mice infected with the knockout strain also showed increased weight loss (Fig. 3B), indicating a more severe illness during the acute phase of the infection.

**Figure 3.**
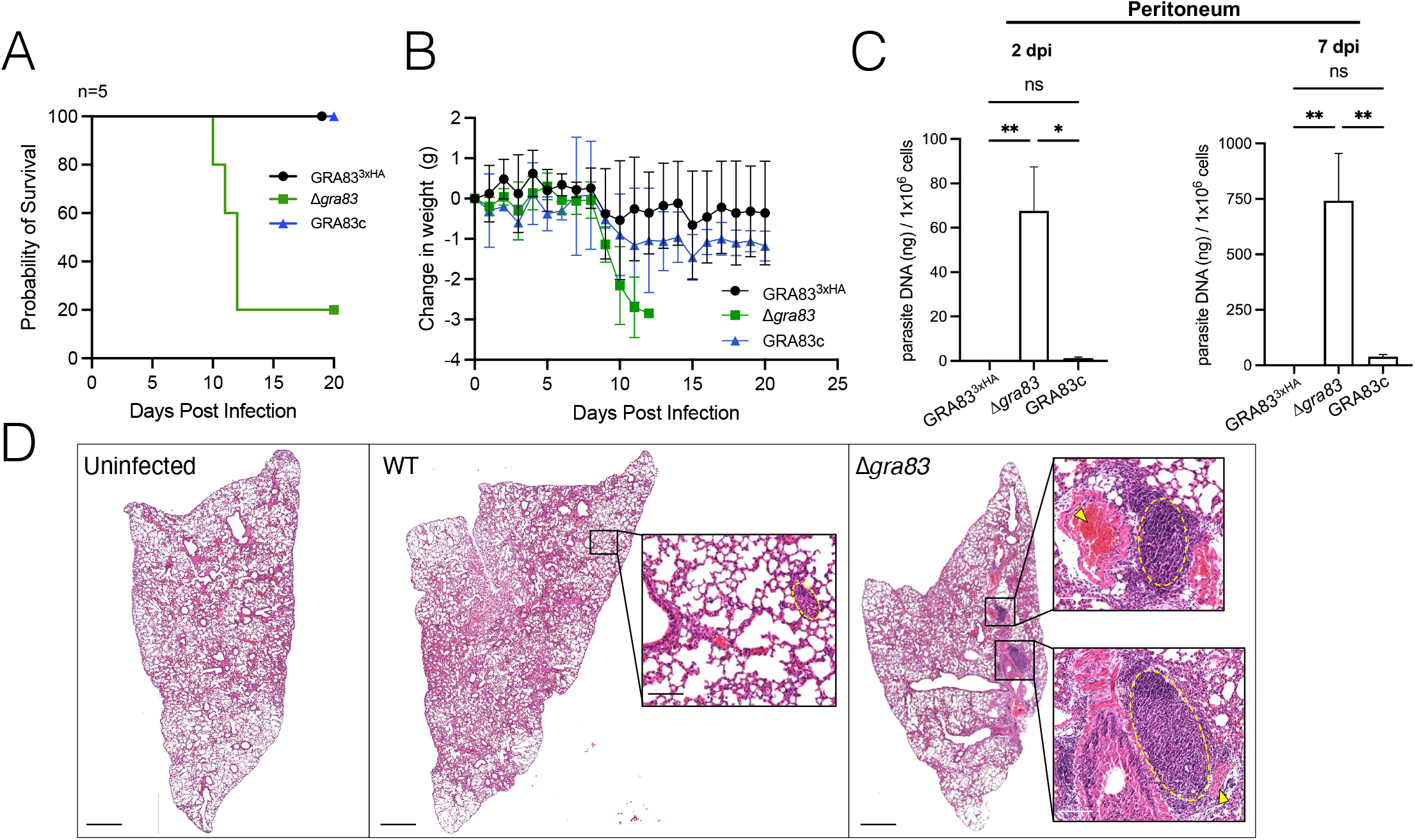
Deletion of *GRA83* increases virulence, parasitemia, and inflammation during the acute infection. A) C57BL/6 mouse survival assay demonstrates the increased virulence of Δ*gra83* parasites (n = 5 per strain). Mice were i.p. infected with 10^3^ parasites each. Differences between groups were compared using Kaplan Meier survival analysis, through log-rank (Mantel-Cox) test (*P <* 0.015). B) Weight change of C57BL/6 mice from the virulence assay shown in panel A. Results are shown as the average weight change per group of infected mice each day. C) C57BL/6 mice infected with 300 parasites were evaluated for parasite burden by qPCR. Peritoneal cells were evaluated for the number of copies of the repetitive 529 bp DNA fragment in *T. gondii.* (n = 5 mice / group, with 2 technical replicates each). Results are expressed as mean ± SEM. Data were analyzed using Kruskal-Wallis one-way ANOVA test; *, *P* < 0.05; **, *P* < 0.005. D) Images of H&E-stained lung sections of uninfected or infected C57BL/6 mice with 300 tachyzoites of the GRA83^3xHA^ and Δ*gra83* strains at 7 dpi. Scale bars = 1 mm. Insets show the magnification of the yellow circled regions highlighting inflammatory infiltrates or bronchus-associated lymphoid tissue (BALT) hyperplasia. Arrows show the migration of mononuclear cells through surrounding blood vessels. Scale bars = 100 µm.

To further explore this phenotype, we assessed parasite burden by qPCR at two and seven dpi (Fig. 3C). In agreement with the enhanced virulence, mice infected with Δ*gra83* parasites had a significantly higher parasite burden in the peritoneum, with levels several fold greater than the GRA83^3xHA^ strain at both time points, and this increased burden was reversed by complementation. We next sought to determine if the increased virulence of the Δ*gra83* parasites led to inflammatory lesions in the nearby lung tissue, one of the preferred targets of parasite replication during the acute phase (45). For mice infected with wild-type control strain, we observed interstitial inflammatory infiltrates throughout the parenchyma of the lungs at 7 dpi, composed mainly of mononuclear cells (Fig. 3D). On the other hand, histological analysis of the lungs of Δ*gra83* infected mice showed compromised tissue integrity due to the extensive interstitial inflammation associated with sites of bronchus-associated lymphoid tissue (BALT) hyperplasia. Together, these data demonstrate that loss of GRA83 results in enhanced growth of the parasites in the acute phase of the infection.

### GRA83 impacts cyst burden and inflammation in the brains of chronically infected mice

To assess whether GRA83 is important for the chronic infection, we first infected CBA/J mice, which produce higher cyst numbers (46), with sublethal doses of tachyzoites to assess weight loss and cyst burden. As observed with the C57BL/6 mice (Fig. 3B), CBA/J mice infected with Δ*gra83* parasites suffered more weight loss than the controls during the acute infection (Fig. 4A), and this weight loss persisted until the end of the study. At 30 dpi, we quantified brain cyst burden and found that Δ*gra83* infection resulted in a significant increase in the cyst burden compared to GRA83^3xHA^ or GRA83c infected mice (Fig. 4B). We did not observe any gross morphological changes of the brain cysts in the mice infected with the Δ*gra83* strain (not shown). To confirm these findings, we similarly challenged C57BL/6 mice with the strains. We again observed an increase in the cyst burden of Δ*gra83* chronically infected mice, as assessed by the presence of *LDH2*-driven GFP^+^ cysts in brain sections (Fig. 4C and D). Although we cannot exclude the possibility that GRA83 plays a direct role in parasite encystment, these data suggest that the deletion of *GRA83* provides the parasite with an advantage during the acute infection which persists and allows *T. gondii* to disseminate to the brain more efficiently, resulting in a higher cyst burden.

**Figure 4.**
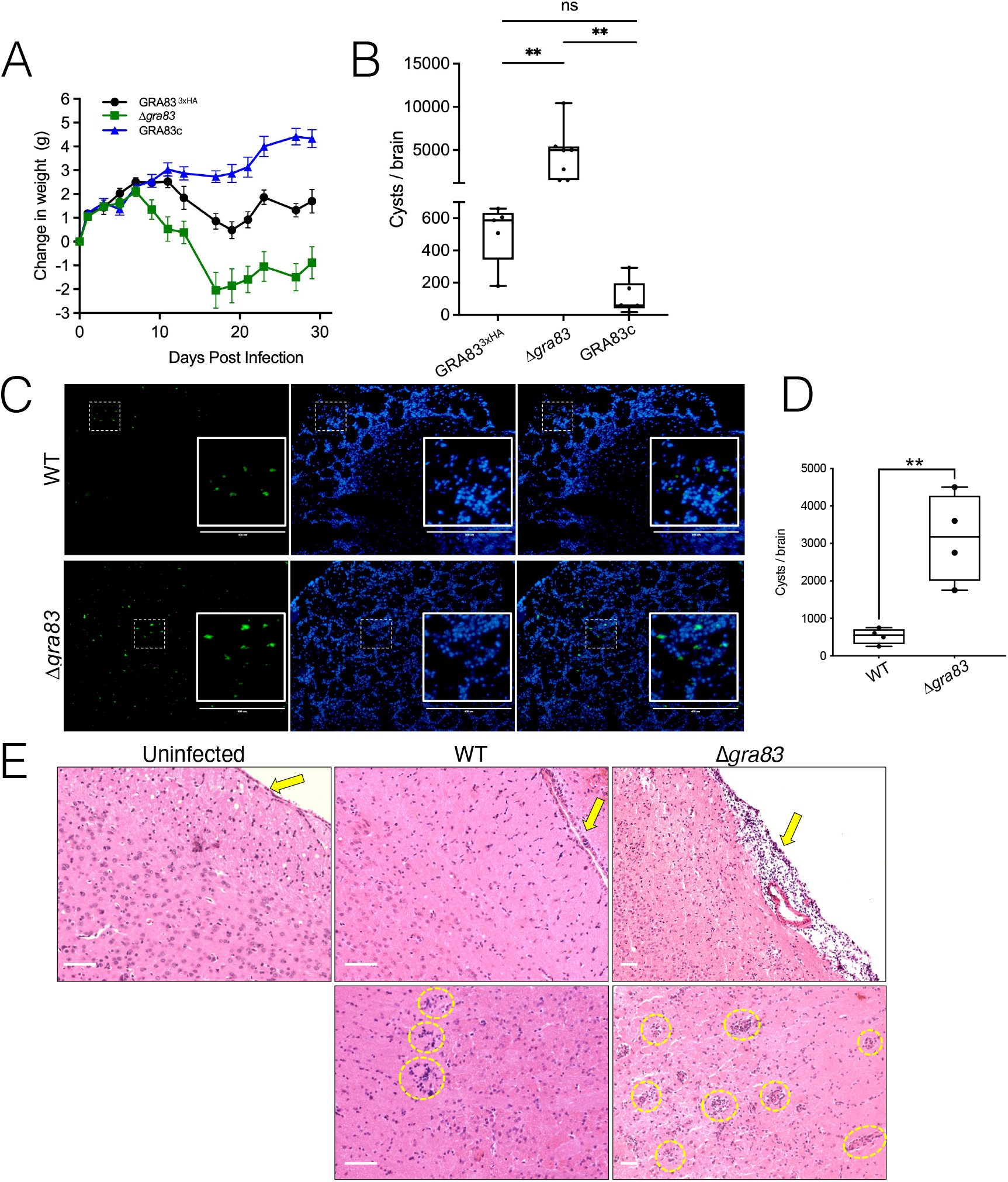
Deletion of *GRA83* increases cyst burden and brain inflammation during the chronic infection. A) Weight change of CBA/J mice infected with 400 parasites of each strain. Results are shown as the average weight change per group of infected mice each day (GRA83^3xHA^: n = 5, Δ*gra83*: n = 7, GRA83c: n = 6). B) Brain cyst burden of CBA/J mice at 30 dpi infected with 400 parasites each of the indicated strains as in panel A. Results are shown as values with min and max whiskers. One-way ANOVA test; **, *P* < 0.01. C) Images of frozen brain sections of infected C57BL/6 mice at 32 dpi with 300 tachyzoites each. Cysts are *LDH2*-GFP^+^ and DAPI labels the cell nuclei. Insets show the magnification of the boxed regions highlighting the cysts. Scale bars = 400 µm. D) Quantification of brain cyst numbers from panel C (n = 4 per group). Results shown as values with min and max whiskers. Two-tailed t-test, **, *P* < 0.05. E) Images of H&E stained brain sections of uninfected or infected C57BL/6 mice with 300 tachyzoites of the GRA83^3xHA^ or Δ*gra83* strains at 30 dpi. Inflammatory infiltrates containing mononuclear cells in the brain parenchyma or around blood vessels are circled in yellow. Arrows point at the meninges of samples analyzed from the different groups showing an intense gathering of mononuclear cells indicating meningitis in mice infected with the Δ*gra83* strain. Scale bars = 50 µm.

We also evaluated inflammation in the cerebral tissue of the chronically infected C57BL/6 mice by histological analysis (Fig. 4E). While few inflammatory infiltrates were seen in sections of the brain parenchyma obtained from mice infected with the wildtype strain, numerous perivascular cuffings of inflammatory mononuclear cells could be observed in brain tissue sections of mice infected with Δ*gra83* strain. Infection with the *Δgra83* parasites was also associated with a severe influx of mononuclear cells to the meninges of those mice, suggestive of initial meningitis.

### GRA83 modulates IL-12/IFN-γ axis *in vitro* and *in vivo*

To further explore the rapid expansion of parasites during the acute infection, we examined whether this phenomenon was due to an imbalance of the IL-12/IFN-γ axis caused by the lack of GRA83. We first measured the levels of IL-12 *in vitro*, using bone marrow derived macrophages (BMDM) infected with the different strains. We observed a significant decrease in the production of IL-12p40 in BMDMs after 18h of infection with the Δ*gra83* strain, compared to the GRA83^3xHA^ or GRA83c infected cells (Fig. 5A).

**Figure 5.**
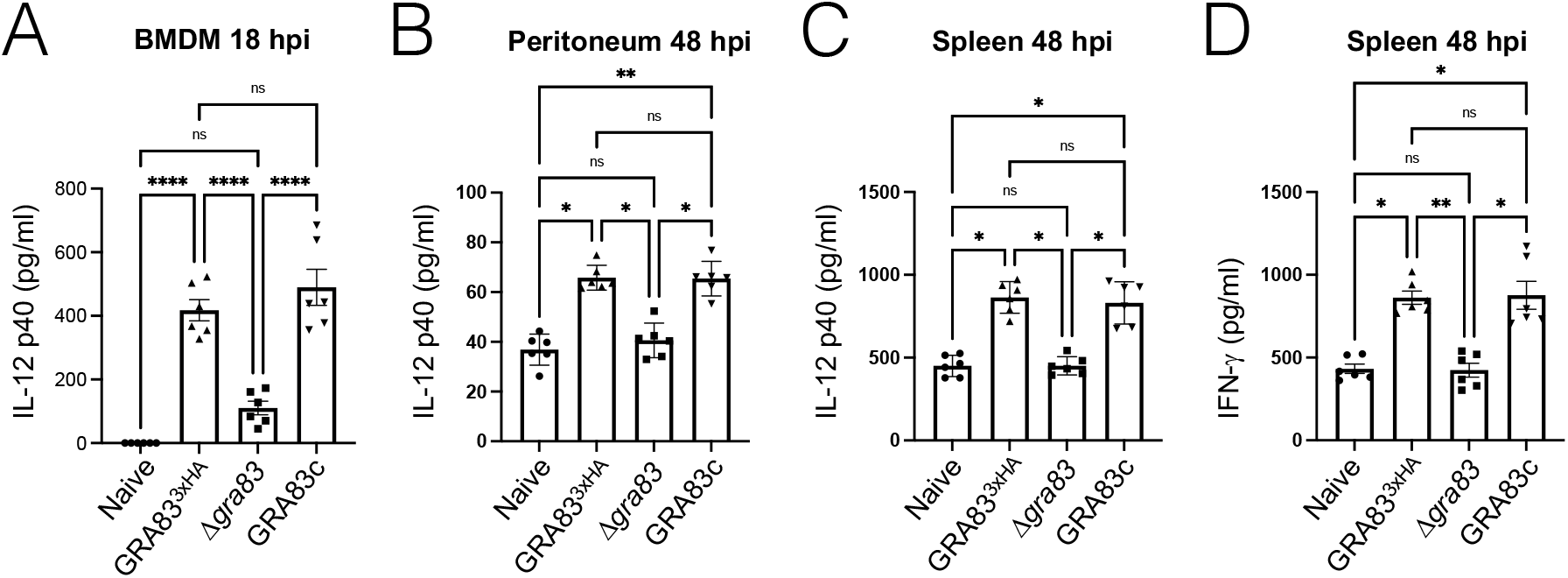
GRA83 impacts Th1 cytokine production *in vitro* and *in vivo*. A) IL-12p40 production from naïve and infected murine BMDMs. Cells were infected with MOI of 0.5 of GRA83^3xHA^, Δ*gra83* and GRA83c strains and supernatants were collected after 18 hpi. Results are expressed as mean ± SEM of 6 biological replicates per group. Data were analyzed using Kruskal-Wallis one-way ANOVA test. (****, *P* < 0.001). B) IL-12p40 production measured from peritoneal exudates of C57BL/6 mice infected with 300 parasites for 48h (n = 6 mice / strain). Values are expressed as mean ± SEM. Data were analyzed using Kruskal-Wallis one-way ANOVA test (*, *P* < 0.001). C, D) IL-12p40 and IFN-γ production measured from spleen homogenates from the mice in B. Values are expressed as mean ± SEM. Data were analyzed using Kruskal-Wallis one-way ANOVA test. (*, *P* < 0.001).

We additionally assessed if IL-12p40 production was altered *in vivo* by determining the cytokine concentration at the initial site of the infection (peritoneal exudates), as well as in spleen homogenates, after 48 hours of infection. We found that IL-12p40 levels were again substantially reduced in both the peritoneal exudates (Fig. 5B) and spleen homogenates (Fig. 5C) of mice infected with the Δ*gra83* parasites compared to the controls. As IL-12 stimulates the production of IFN-y, which is the crucial mediator responsible for the clearance of *T. gondii,* we also measured the concentration of this cytokine in the spleen samples. Accordingly, we found that IFN-y production was also dramatically reduced in Δ*gra83* infected mice compared to the controls (Fig. 5D). Together, these results suggest that the absence of GRA83 in *T. gondii* disrupts the critical pro-inflammatory IL-12/IFN-γ axis in mice, allowing uncontrolled parasite replication and severe illness for the host.

### NF-κB nuclear translocation is reduced in Δ*gra83* parasitized cells

NF-κB is a central regulator of IL-12 and the subsequent IFN-γ responses to *T. gondii* infections (19, 47). To determine whether disruption of *GRA83* affects the nuclear translocation of NF-κB, we measured nuclear fluorescence intensity of the NF-κB p65 subunit in HFFs. We observed that infection with Δ*gra83* parasites resulted in significant decrease in p65 nuclear translocation compared to the GRA83^3xHA^ or GRA83c strains (Fig. 6A-E). GRA15 expressed by Type II strains of *T. gondii* activates nuclear translocation of NF-κB in infected HFFs through its direct binding to TRAF2 and TRAF6 (31, 32). To assess the role of GRA15 in the *GRA83* knockout strain, we first tagged *GRA15* in Δ*gra83* parasites which did not show any gross changes in localization or levels of expression of the protein (Fig. S2A and B). We then generated a double knockout of *GRA83* and *GRA15* (Δ*gra83/Δgra15)* to determine whether deletion of both GRA proteins leads to an additive reduction in NF-κB translocation (Fig. S3A and B). However, we found no difference in NF-κB translocation between the Δ*gra83* and Δ*gra83/Δgra15* parasites (Fig. 6E-G). Thus, GRA83 – alongside GRA15 – is important for NF-κB p65 translocation to the nucleus, which leads to the production of IL-12 and, consequently, triggers IFN-γ in an *in vivo* setting. The fact that the double knockout does not appear to have an additive effect on NF-κB activation suggests that these GRAs converge to regulate this important transcription factor involved in regulating the host’s response to the infection.

**Figure 6.**
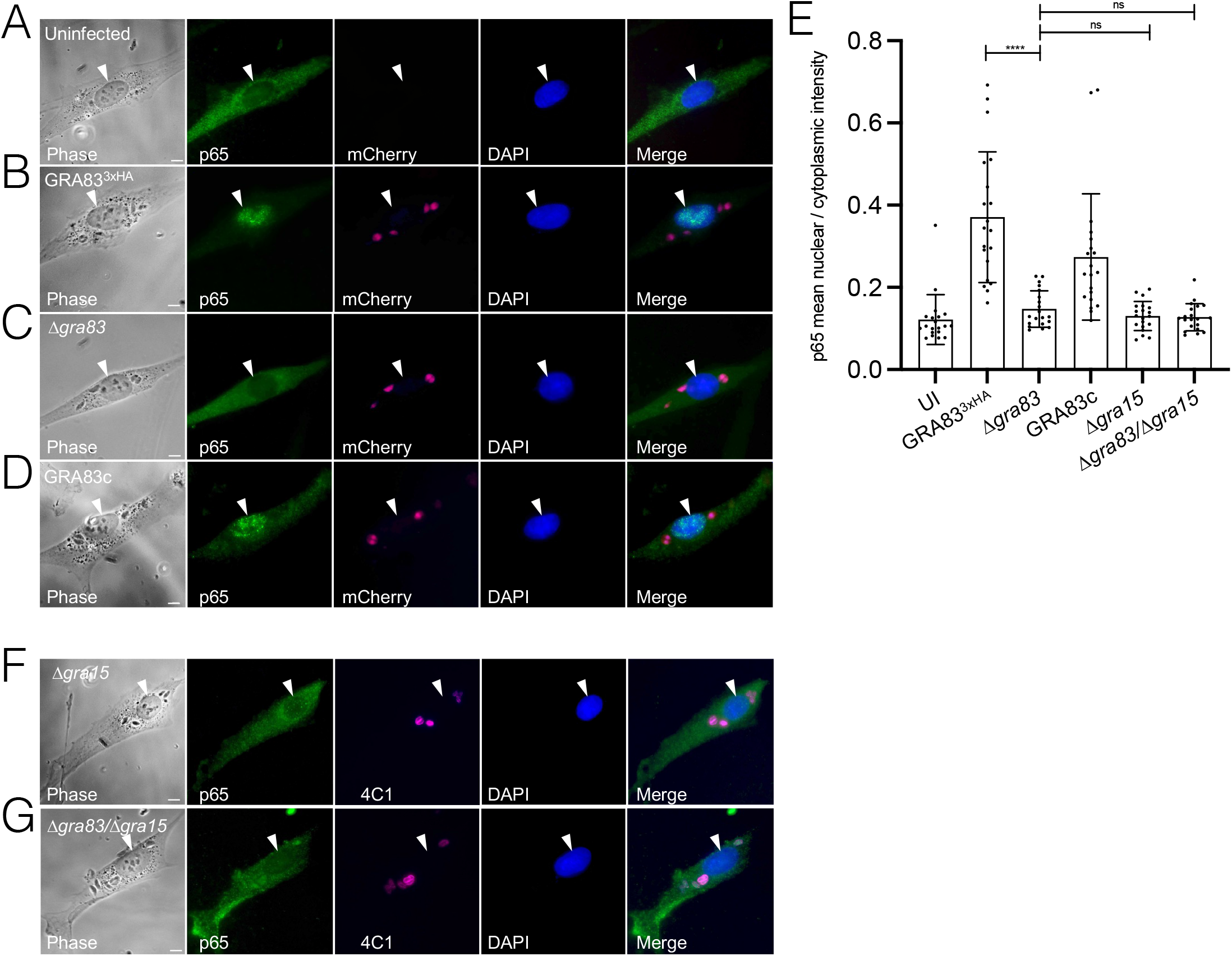
Infection with Δ*gra83* parasites results in reduced NF-κB p65 nuclear translocation in HFFs. A-D) IFA of HFFs infected with *T. gondii* strains for 18 hpi, fixed, and stained with rabbit anti-p65 (green) and DAPI (blue). GRA83^3xHA^, Δ*gra83*, and GRA83c parasites expressed cytosolic mCherry (magenta) to visualize infected cells. F-G) IFA of HFFs infected with Δ*gra15* or Δ*gra83*/Δ*gra15* parasites 18 hpi. Magenta: mouse anti-parasite periphery (mAb 4C1); Green: rabbit anti-p65; Blue: DAPI. E) The ratio of mean p65 nuclear intensity to p65 cytoplasmic intensity was quantitated in 20 HFFs for each strain. Asterisks indicate significantly higher levels of nuclear p65 compared with uninfected cells (****, *P* < 0.0001, ordinary one-way ANOVA test). Each arrowhead points to the host cell nucleus. Scale bars = 10 µm.

### Proximity labelling identifies putative GRA83 parasite interactors

To identify candidate GRA83 interacting proteins from *T. gondii*, we used *in vivo* biotinylation with GRA83 as the bait. To do this, we endogenously tagged GRA83 with TurboID-3xHA (Fig. 7A) and showed that the fusion protein properly targeted to the PV, similar to that seen for the 3xHA tagged protein (Fig. 7B). Addition of biotin to the media showed robust streptavidin labeling in the vacuole overlapping with the fusion protein, demonstrating that the fusion protein was active (Fig. 7C). We then conducted large scale proximity labeling experiments, purified the biotinylated proteins, and identified them by mass spectrometry (Fig. 7D and S4). As expected, GRA83 was the top *T. gondii* hit in the dataset, likely due to self-biotinylation of the bait protein. In addition, an array of known GRA proteins involved with modulating the host during *T. gondii* infection were identified in the top 25 hits. This included GRA15, suggesting this may be a GRA83 partner in the PV. We attempted to assess GRA15/83 interaction in dual tagged parasites by co-immunoprecipitation and western blotting, but we were unable to observe any significant interaction (not shown). Another top hit in the proximity labeling was GRA57, which is also present in the PVM and has been reported to be involved with IFN-γ response (33, 48). GRA28 was also identified, which localizes to the host cell nucleus in a MYR1- and ASP5-dependent manner and facilitates parasite dissemination in hosts and drive migration of infected macrophage (28, 49). In agreement with several of these GRAs that are secreted into the host cell, MYR1, a component of the PVM translocon, was also ranked in the top 25 hits.

**Figure 7.**
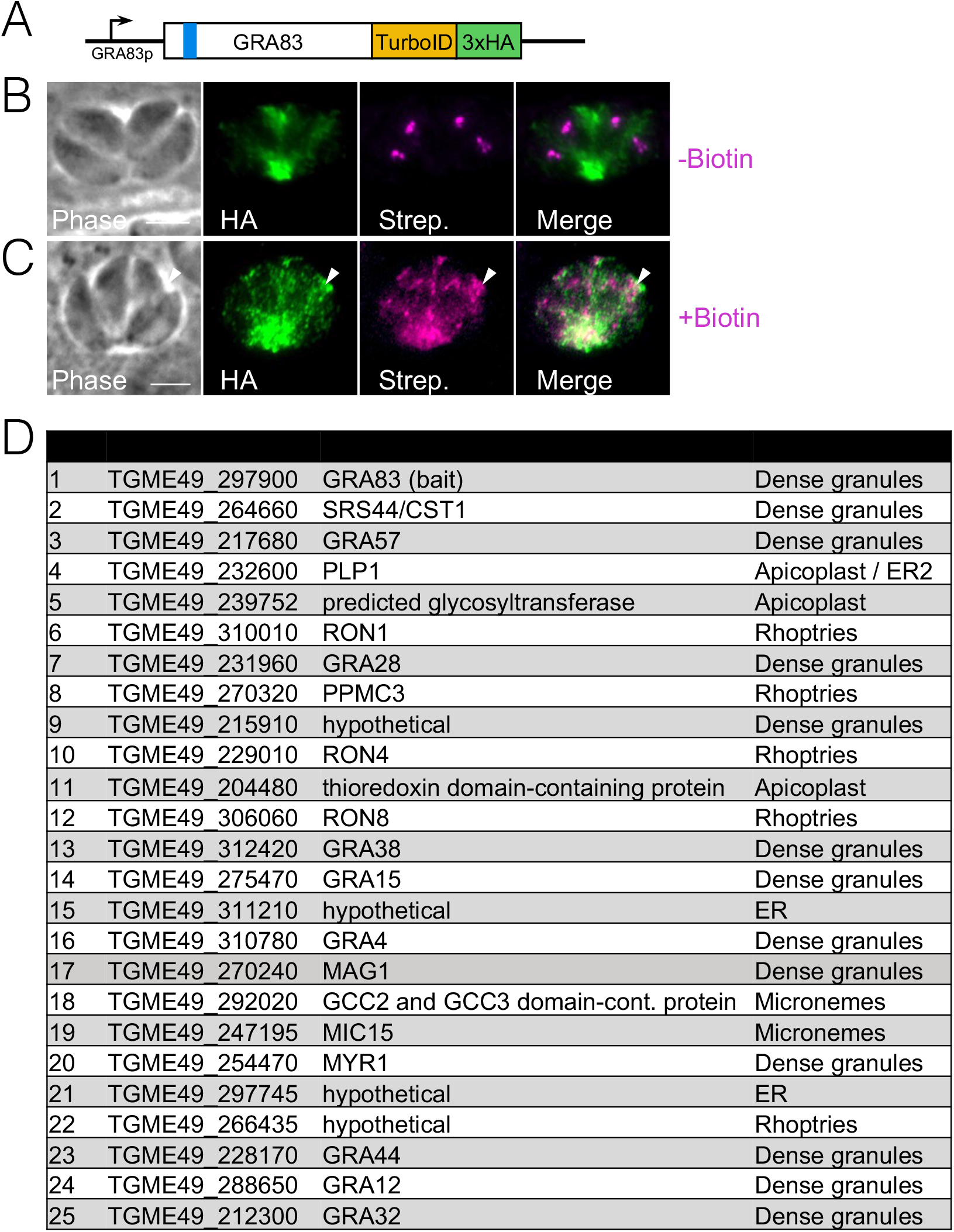
GRA83^TurboID-3xHA^ localizes to the PV and biotinylates target proteins in the PV. A) Diagram showing the C-terminally tagged GRA83 with TurboID-3xHA. B) IFA of GRA83^TurboID-3xHA^ without biotin supplemented to the media. The fusion protein appropriately localizes to the tachyzoite vacuole. Endogenously biotinylated proteins are detected at the apicoplast of each parasite. Magenta: streptavidin; Green: mouse anti-HA. C) IFA of the GRA83^TurboID-3xHA^ expressing parasites grown for 24h in media supplemented with biotin. Streptavidin labels the PV where the fusion protein accumulates (arrowheads). Magenta: streptavidin; Green: mouse anti-HA. Scale bar = 5 µm. D) Table of the top hits from the GRA83-TurboID *in vivo* biotinylation experiment. Hits were ranked based on spectral count number and absence in the control group.

## Discussion

In this work, we characterize the function of the secreted *Toxoplasma* protein GRA83. GRA83 is larger than most GRAs with a predicted mass of ∼83 kDa, but similar to other GRAs it lacks identifiable domains that allude to its function. The protein is largely unstructured (∼40% disordered), but this is on the lower side compared with other host cell localizing GRAs (e.g. GRAs 6, 16, 24, TgIST) that range from 60 - 80% disordered (25, 29, 50, 51). We show that GRA83 is secreted into the vacuole in both tachyzoites and bradyzoites, suggesting that it functions in both lifecycle stages. While this manuscript was in preparation, another group also demonstrated that the epitope tagged protein is secreted into the vacuole in tachyzoites (52). Together with the hyperLOPIT prediction (34), which tends to be reliable for dense granule proteins, these results confirm that GRA83 is indeed a secreted GRA protein.

We determined that the N-terminal portion of the protein containing a hydrophobic region is sufficient for secretion, although some of the truncated protein appears arrested in the parasite’s secretory pathway. This may be due to misfolding of our fusions to the smOLLAS tag which adopts a GFP-like fold (43), whereas most GRAs are unstructured, a feature that may aid in traversing the secretory pathway. Similar N-terminal hydrophobic domains have been identified in other *T. gondii* GRA proteins including GRA6, TgNSM, and GRA15 where they may also serve as non-canonical secretory signals (30, 31, 50). As these signals are not believed to be cleaved, the hydrophobic domain may also play another role (e.g. membrane association) upon delivery to its destination. While the level of membrane association or topology of GRA83 is unknown, it may be similar to GRA6 which is believed to be tethered to the cytoplasmic face of the PVM via its N-terminal hydrophobic domain (50). Alternatively, GRA83’s function may occur entirely within the vacuole where it may reside as a membrane associated or soluble protein. Interestingly, similar internal hydrophobic regions are also found in several secreted proteins in *Plasmodium spp.*, where they have recently been shown to play a direct role in ER insertion and delivery of proteins into the secretory pathway (53).

Similar to many *Toxoplasma* secreted effector proteins that modulate host functions, GRA83 can be disrupted and the Δ*gra83* parasites grow normally in HFFs *in vitro*. This agrees with the phenotype score of 0.94 in the genome-wide CRISPR screen of type I (RH strain) parasites (54). However, infection with *Δgra83* parasites leads to a substantial increase in weight loss, parasitemia, and overall virulence in infected mice. Assessing the immune response to Δ*gra83* parasites revealed that this protein is able to induce IL-12p40 production. This triggers the assembly of critical IFN-γ dependent effector responses that will actively inhibit parasite replication, dampening the potential to cause extensive lesions in several tissues (9). The higher number of brain cysts in the chronic infection may simply be due to the higher parasitemia in the acute infection, but this may also reflect a similar role for GRA83 continuing to regulate the immune response in bradyzoites. This latter possibility is supported by data showing that GRA83 is upregulated in bradyzoites (42). It would be interesting to test whether GRA83 driven from a tachyzoite-specific promoter still resulted in higher cyst numbers in the chronic infection.

The dampened immune response in Δ*gra83* infected mice correlates with a reduction in NF-κB activation, which is known to control IL-12 mediated IFN-γ production (9, 30, 55). NF-κB has been shown to be primarily activated by the polymorphic GRA15 protein from the type II background strain of *T. gondii,* although GRA7 and GRA14 also have been reported to play less substantial roles in activation (30, 56). Loss of GRA15 results in M2 macrophage polarization through decreased NF-κB activation, a reduction in IL-12 and IFN-γ production *in vivo*, and increased parasite burden, a pattern similar to what we observe for GRA83 (30, 55, 57). However, reports on the virulence of Δ*gra15* parasites appear to disagree on the increased lethality induced by the parasite in mice, a phenotype that we have confirmed in repeated experiments using Δ*gra83* parasites (30, 57–59). The mechanism by which GRA15 limits pathogen growth in IFN-γ stimulated human and murine fibroblasts is through direct interaction with the host ubiquitin ligases TRAF2 and TRAF6 at the vacuolar membrane, which enhances ubiquitin mediated disruption of the vacuole (31, 32). Human monocytes also rely exclusively on GRA15 to produce the acute inflammatory IL-1β through the activation of the proteolytic inflammasome complex (60). In addition, GRA15 has been shown to activate the cGAS/STING pathway due to its localization in the host cell ER after secretion, again limiting parasite expansion and favoring the chronic infection (57). While we show that GRA83 regulates NF-κB in a similar manner to that observed for GRA15, it is unlikely to directly bind TRAF2 or TRAF6, as the protein lacks the binding sites for these factors that are present in GRA15 (TRAF2: (P/S/A/T)X(Q/E)E or TRAF6: (PXEXXZ)) (32). Determining the precise mechanism by which GRA83 regulates NF-κB nuclear translocation will provide new insight into how it regulates this central control element of infection.

We were unable to observe an additive effect on the reduction of NF-κB activation from the double knockout of *GRA15* and *GRA83*. This is likely due to the convergence of both secreted proteins in the same host pathway for modulating activation. One possibility is that GRA83 and GRA15 interact and regulate each other’s function, either within the vacuole or upon its destination at the PVM. Our proximity labeling with GRA83 as bait showed that GRA15 was highly ranked in the dataset, supporting this possibility. While we did not see GRA83 and GRA15 interaction by co-immunoprecipitation, this could be due to a weak interaction between the proteins or one that is unstable in detergent lysates.

While most effector proteins are aimed at interfering with the immune response to avoid pathogen clearance, it is intriguing that *T. gondii* expresses GRA15 and GRA83 that are able to limit the infection via stimulating NF-κB (30). Particularly when the parasite also secretes effectors such as TEEGR and ROP38 that inhibit NF-κB mediated responses (61, 62). This is likely important for regulating this pathway at different timepoints in the infection, different host cell types, or for balancing the infection in a variety of hosts. While *N. caninum* has a GRA83 orthologue, it lacks GRA15 as well as several other effectors that target host survival or immune evasion (e.g. GRA24, GRA28, TgIST1, TEEGR, among others) (23). *N. caninum* also does not significantly activate NF-κB (63), suggesting that GRA83 may play a role in enabling GRA15 to bind to TRAF2/6 and consequently translocate NF-κB the nucleus, or that a second function for GRA83 in both organisms has yet to be discovered. Recently developed tools for genetic manipulation of *N. caninum* will aid in the already proven approach of comparing these pathogens using heterologous expression or knockouts of various effectors to assess how individual or groups or effectors are able to mediate infection of these pathogens in their various mammalian hosts (26, 64).

In summary, GRA83 is a secreted dense granule protein that is expressed in different life stages of *T. gondii*. It functions, by a yet unknown molecular mechanism, to regulate NF-κB translocation as well as IL-12 and IFN-γ production that help limit parasite growth *in vivo*. Together, GRA83 serves as a “pro-host” effector that counterbalances an array of “pro-parasite” effectors which collectively fine tune the host’s immune response, likely to ensure host survival and ultimately enable transmission.

## Methods

### Host cells and parasite culture

Parental PruΔ*ku80*Δ*hxgprt* (Type II, strain Pruniaud) and modified strains of *T. gondii* were maintained on confluent monolayers of human foreskin fibroblast (HFF) (BJ, ATCC, Manassas, VA) and cultured as previously described (33). Constructs containing selectable markers were selected using 1 µM pyrimethamine (for dihydrofolate reductase-thymidylate synthase [DHFRTS]) or 50 µg/mL mycophenolic acid-xanthine (for hypoxanthine-xanthine-guanine phosphoribosyl transferase [HXGPRT]) (65, 66). Homologous recombination to the *UPRT* locus was negatively selected using 5 µM 5-fluorodeoxyuridine (FUDR) (44).

### Antibodies

The antibodies used in this study were mouse monoclonal antibody (mAb) HA.11 (Biolegend; 901515), rabbit polyclonal antibody (pAb) anti-HA (Invitrogen; PI715500), rat mAb anti-OLLAS (67), mouse mAb anti-4C1 (68), rabbit pAb anti-IMC6 (69), mouse pAb anti-RON5c (70), mouse pAb anti-GRA7 (12B6) (41), mouse pAb anti-GRA14 (41), and rabbit anti-NF-κB p65 (Cell Signaling Technology; D14E12). Biotinylated proteins were detected with streptavidin Alexa Fluor 546 conjugate (Molecular Probes; S-11225).

### Epitope tagging and gene disruption

For C-terminal endogenous tagging, a pU6-Univeral plasmid containing a protospacer against the 3’ untranslated region (UTR) of the gene of interest (∼150 bp downstream of the stop codon) was generated, as previously described (71). A homology-directed repair (HDR) template was PCR amplified using the Δ*ku80*-dependent LIC vector p3xHA.LIC-DHFR, smOLLAS.LIC-DHFR, or TurboID-3xHA-DHFR, which contained the epitope tag, 3’ UTR, and a selection cassette (43). The HDR templates include 40 bp of homology immediately upstream of the stop codon or 40 bp of homology within the 3’ UTR downstream of the CRISPR/Cas9 cut site. Primers that were used for the pU6-Universal plasmid as well as the HDR template are listed in Table S1 as P1-P10.

For gene deletions, the protospacer was designed to target the coding region of the target gene and ligated into the pU6-Universal plasmid and prepared similarly to the endogenous tagging constructs (Table S1, P11-12 and P15-16). The HDR template included 40 bp of homology immediately upstream of the start codon or 40 bp of homology downstream of the region used for homologous recombination for endogenous tagging (Table S1, P13-14 and P17-18). The HDR template was PCR amplified from a pJET vector containing the *HXGPRT* drug marker driven by the NcGRA7 promoter.

For all tagging and gene deletions, the HDR template was amplified in 400 μL, purified by phenol-chloroform extraction, ethanol precipitated, and electroporated into appropriate parasite, along with 50 μg of the pU6-Universal plasmid. Transfected parasites were allowed to invade a confluent monolayer of HFFs overnight, and the appropriate selection drug was subsequently applied. Successful tagging or deletion was confirmed by IFA, and clonal lines of tagged parasites were obtained through limiting dilution. Knockout clones were verified by PCR to examine for the absence of the gene of interest and presence of the *HXGPRT* cassette at the predicted recombination site (Table S1, P19-25).

To generate a Δ*gra83*/Δ*gra15* mutant, we first removed the *HXGPRT* cassette from the Δ*gra83* strain by transfecting in linearized NcGRA7p-pJET plasmid and pU6-Universal plasmid containing a protospacer targeting the NcGRA7 promoter. We selected *HXGPRT*-negative clones using 6-thioxanthine (72). We next C-terminally tagged *GRA15* with the 3xHA DHFR in the parasites and subsequently deleted *GRA15* using the methods described above.

### Complementation of GRA83

GRA83-3xHA was re-introduced into the genome of the Δ*gra83* strain in the *UPRT* locus. Primers were designed to PCR amplify the GRA83 native promoter from genomic *T. gondii* DNA of PruΔ*ku80*Δ*hxgprt* (Table S1, P26-P27) and amplify the *GRA83* gene from a PruΔ*ku80*Δ*hxgprt* cDNA template (Table S1, P28-29). This was cloned into the pUPRT knockout vector, resulting in the pUPRTKO-GRA83p-GRA83-3xHA. This complementation vector was linearized with PsiI-v2 (NEB), transfected into Δ*gra83* parasites, and selected with FUDR. Clones were selected by limiting dilution, examined by IFA (anti-HA antibody), and a positive clone was designated GRA83c.

### mCherry expressing GRA83^3xHA^, Δ*gra83*, and GRA83c *T. gondii*

100 ug of either pTub.mCherry.3xHA.UPRT or pTub.mCherry.3xHA.DHFR plasmids were linearized using PsiI-HF. GRA83^3xHA^ or Δ*gra83* parasites were transfected with the linearized pTub.mCherry.3xHA.UPRT plasmid and PU6 plasmid containing gRNA targeting the *UPRT* locus and selected with FUDR. GRA83c parasites were transfected with linearized pTub.mCherry.3xHA.DHFR and selected with pyrimethamine. After selection, parasites were cloned by limiting dilution and cytosolic mCherry expression was confirmed by IFA.

### Immunofluorescence assay and western blot

For IFAs, glass coverslips of confluent HFFs were infected with *T. gondii* for approximately 24-34 hours. The coverslips were then fixed in 3.7% formaldehyde in PBS and processed for immunofluorescence as previously described (33). Primary antibodies were detected by species-specific secondary antibodies conjugated to Alexa Fluor 488/594 (ThermoFisher). Coverslips were mounted onto microscope slides with Vectashield (Vector Labs, Burlingame, CA), or Vectashield with DAPI mounting media and the fluorescence was observed using a Zeiss Axio Imager.Z1 microscope. Images were processed with Zeiss Zen 2.3 software (Zeiss).

For the IFAs intended for determining NF-κB p65 translocation, HFFs were infected with *T. gondii* strains for 18 h, fixed, and stained, as described above. Vectashield with DAPI was used to mount coverslips onto the microscope slides. Nuclear p65 translocation was determined by calculating the ratio of the mean nuclear p65 intensity divided by the mean cytoplasmic p65 intensity, and graphs were made using Prism GraphPad. This was quantitated in 20 infected HFF cells for each strain.

For western blot, intracellular parasites were lysed in 1x Laemmli sample buffer with 100 mM DTT and boiled at 100°C for 10 min. Lysates were resolved by SDS-PAGE and transferred to nitrocellulose membranes overnight at 4°C. Proteins were detected with the appropriate primary antibody and corresponding secondary antibody conjugated to horseradish peroxidase. Chemiluminescence was induced using the Supersignal West Pico substrate (Pierce) and imaged on a ChemiDoc XRS+ (Bio-Rad, Hercules, CA).

### *In vitro* conversion of *T. gondii* tachyzoites to bradyzoites

HFFs were grown on coverslips for seven days and infected with GRA83^3xHA^ parasites for 3h. The media was changed to bradyzoite “switch” media (DMEM with 1% FBS, 50mM HEPES, 1% PSG, pH 8.1) and the cultures were incubated at 37°C in ambient air (0.05% CO_2_) for 3 days to induce conversion to bradyzoites, as previously described (33, 73). The coverslips were then fixed for IFA analysis as above.

### Plaque assay

Six-well plates were seeded with confluent monolayers of HFFs and infected with approximately 250 parasites and allowed to form plaques for eight days. The monolayers were then fixed in 100% methanol for 3 minutes, washed with PBS, and stained with crystal violet for visualization. The areas of at least 50 plaques per strain were measured using Zen software (Zeiss), and experiments were done in triplicates. Plaque means were compared using one-way ANOVA test. Data was graphed with GraphPad Prism (used to make all graphs).

### Mouse virulence assay

GRA83^3xHA^, Δ*gra83*, or GRA83c parasites were collected from infected NIH-3T3 monolayers and resuspended in RPMI 1640 medium (Thermo Fisher Scientific) prior to intraperitoneal injection into female C57BL/6 mice (6-8 weeks of age), with 10^3^ tachyzoites. A sample of each inoculum was stored for later confirmation by qPCR. Mice were monitored for symptoms of infection, weight loss, and survival for 20 days.

### Determination of parasite burden

The parasite burden was determined in C57BL/6 mice infected with GRA83^3xHA^, Δ*gra83*, and GRA83c tachyzoites. Peritoneal cells were submitted to a real-time qPCR using SYBR green detection system (PowerUp SYBR Green Master Mix, Thermo). Genomic DNA was extracted by salting out. DNA concentrations were quantified by spectrophotometer (260/280 ratio; Nanodrop Lite, Thermo Scientific) and adjusted to 40 ng/µL with sterile DNAse free water. The reaction to determine parasite load was performed in a Real-time PCR thermal cycler (StepOne Plus, Thermo Scientific, USA) and parasite genomic DNA was quantified by interpolation from a standard curve with known amounts of DNA extracted from *T. gondii* tachyzoites that were included in each analysis. Primers used to examine the repetitive 529 bp DNA fragment in *T. gondii* are included in Table S1 as P30-31.

### Histological analysis

Lung and brain samples of *T. gondii* infected C57BL/6 mice were collected and fixed in 10% buffered formalin at room temperature for 24 h and stored in alcohol 70% until the paraffin inclusion process. After inclusion, the organs were sliced (5 mm thick) and deposited on microscopic slides, subsequently stained with hematoxylin and eosin for evaluation of inflammatory infiltrates and tissue damage as previously described (74). The stained sections were photographed using an automated slide scanner microscope (Aperio ScanScope AT, Leica).

### Mouse brain cyst quantitation

Intracellular GRA83^3xHA^, Δ*gra83,* and GRA83c parasites were mechanically liberated from infected HFF monolayers and resuspended in Opti-MEM prior to intraperitoneal injection into groups of female CBA/J mice each. 400 tachyzoites per mouse of each strain were injected. Injected parasites were confirmed to be live and viable parasites by plaque assays with HFF monolayers. The mice were monitored for symptoms of infection and weight loss 30 days after injection and then sacrificed. Mouse brains were collected and *LDH2-*GFP+ tissue cysts were quantified, as previously described (33).

For the C57BL/6 strain, mice were infected with 300 tachyzoites and monitored for 32 days for clinical signs. Mice were then euthanized and had the brain removed and divided in half: one hemisphere was homogenized and examined by phase microscopy for tissue cysts; the other hemisphere was placed in embedding medium (Tissue-Tek O.C.T. Compound, Sakura Finetek) and quickly frozen in liquid nitrogen. The tissues were submitted to cryosectioning and counterstained with DAPI. The slides were read in a fluorescent microscope (EvosFL, Thermo) and tissues cysts were identified by the expression of *LDH2*-GFP+.

### Cytokine measurements

C57BL/6 BMDMs were differentiated and stimulated as previously described (75). Briefly, the cells were seeded (2 x 10^5^ per well) in 96-well plates and left to adhere overnight at 37°C in 5% CO_2_. Cells were infected with freshly lysed GRA83^3xHA^, Δ*gra83,* and GRA83c tachyzoites at MOI of 0.5, and supernatants (200 µl) were collected 18 h after infection and stored at −80°C. For *in vivo* assays, C57BL/6 mice were euthanized after 48h of infection and the peritoneal cavity was washed with 1 ml of PBS, spun at 400 x g for 10 min to pellet the cells. The supernatant was collected and stored at −80°C. Spleens were also collected at 48 h, mechanically disrupted by a tissue homogenizer (Ika) in 1 ml of PBS with protease inhibitor cocktail (Roche) and spun at 10,000 x g for 10 min to pellet debris. Supernatants were collected and stored at −80°C. IL-12p40 and IFN-γ levels were determined using commercially available ELISA kits (BD Biosciences) according to the manufacturer’s instructions.

### Affinity capture of biotinylated proteins

HFF monolayers were infected with parasites expressing the GRA83^TurboID-3xHA^ or the parental line. Upon infection, 150 μM biotin was supplemented in the media for 30 h. Intracellular parasites were collected, washed in PBS, and lysed in radioimmunoprecipitation assay (RIPA) buffer (50mM Tris [pH 7.5], 150 mM NaCl, 0.1% SDS, 0.5% sodium deoxycholate, 1% NP-40) supplemented with Complete Protease Inhibitor cocktail (Roche) for 30 min on ice. Lysates were centrifuged for 15 min at 14,000 × g to pellet insoluble material and the supernatant was incubated with Streptavidin Plus UltraLink Resin (Pierce) at 4°C overnight under gentle agitation. Beads were collected and washed five times in RIPA buffer, followed by three washes in 8 M urea buffer (50 mM Tris-HCl pH 7.4, 150 mM NaCl). Ten percent of each sample was boiled in Laemmli sample buffer, and eluted proteins were analyzed by western blotting by streptavidin-HRP while the remainder was used for mass spectrometry.

### Mass spectrometry of biotinylated proteins

The proteins bound to streptavidin beads were reduced and alkylated via sequential 20-minute incubations of 5mM TCEP 10 mM iodoacetamide at room temperature in the dark while being mixed at 1200 rpm an Eppendorf thermomixer. Proteins were then digested by the addition of 0.1μg Lys-C (FUJIFILM Wako Pure Chemical Corporation, 125-05061) and 0.8 μg Trypsin (Thermo Scientific, 90057) while shaking 37°C overnight. The digestions were quenched via addition of formic acid to a final concentration of 5% by volume. Each sample was desalted via C18 tips (Thermo Scientific, 87784) and then resuspended in 15 μL of 5% formic acid before analysis by LC-MS/MS.

Peptide samples were separated on a 75 uM ID, 25 cm C18 column packed with 1.9 μM C18 particles (Dr. Maisch GmbH) using on a 140-minute gradient of increasing acetonitrile and eluted directly into a Thermo Orbitrap Fusion Lumos instrument where MS/MS spectra were acquired by Data Dependent Acquisition (DDA). Data analysis was performed using the ProLuCID and DTASelect2 algorithms (76–78) as implemented in the Integrated Proteomics Pipeline - IP2 (Integrated Proteomics Applications, Inc., San Diego, CA) against the *T. gondii* ME49 reference proteome from ToxoDB and the human reference proteome from Uniprot. Protein and peptide identifications were filtered using DTASelect and required a minimum of two unique peptides per protein and a spectrum-level false positive rate of less than 1% as estimated by a decoy database strategy. Candidates were ranked by spectral counts comparing GRA83^TurboID^ versus control samples.

### Animal experimentation ethics statement

Specific details of our protocol were approved by the UCLA Institutional Animal Care and Use Committee (protocol: 2004-005). Experiments conducted at the Institute of Biomedical Sciences, Federal University of Uberlândia were approved by the institution’s animal research ethics committee (Comitê de Etica na Utilização de Animais da Universidade Federal de Uberlândia—CEUA/UFU), under protocol number 109/16, and were carried out in accordance with the recommendations in the International Guiding Principles for Biomedical Research Involving Animals of the International Council for Laboratory Animal Science (ICLAS), countersigned by the Brazilian National Council for the Control of Animal Experimentation (Conselho Nacional de Controle de Experimentação Animal, CONCEA). REBIR/UFU is accredited by the National Commissions in Animal Experimentation (CONCEA, CIAEP 01.0105.2014) and Biosecurity (CTNBio, CQB 163/02).

For the chronic infection model, these mice were sacrificed at the conclusion of the experiment at 30 days post-infection. Euthanasia was done per AVMA guidelines—using slow (20-30% per minute) displacement of chamber air with compressed CO_2_. This is followed by confirmatory method of cervical dislocation.

## Acknowledgments

The authors thank Drs. Upinder Singh, John Boothroyd, and David Bzik for sharing the Pruniaud Δ*ku80*Δ*hxgprt* strain that contains GFP driven from the *LDH2* promoter. The authors also thank members of the Bradley and Mineo laboratories for their reading and comments of the manuscript. The funders of the study had no role in study design, data collection and interpretation, or the decision to submit the work for publication. This work was supported by NIH grants AI125106 and AI123360 to P.J.B. and GM089778 to J.A.W.; A.C.T. was supported by the Ruth L. Kirschstein National Research Service Award AI007323 and the UCLA Molecular Biology Institute (MBI) Whitcome Fellowship. This work was also supported by Brazilian funding agencies (FAPEMIG - CVZ-PPM-00547-17, CVZ-APQ-01313-14, RED-00313-16). T.W.P.M. was individually supported by Brazilian National Council for Scientific and Technological Development (313761/2020-5) and CAPES (88881.173280/2018-01) fellowships. C.M.M. and A.P.N.G. were also supported by CAPES fellowships.

---

**Fig. S1.**
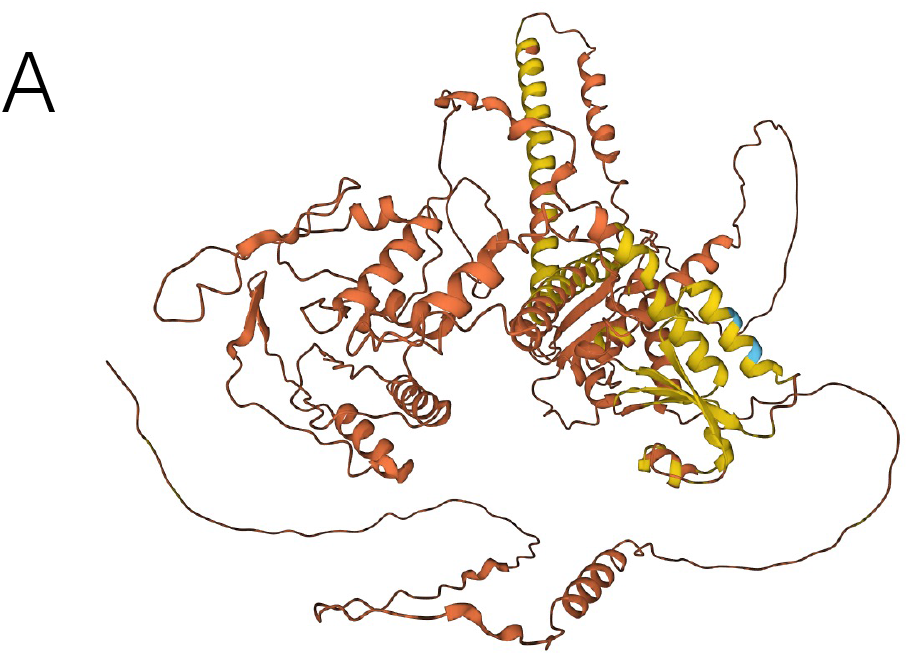
Alphafold provides a low confidence structure of GRA83. A) AlphaFold structure prediction generates an overall low confidence model for GRA83. Blue segments represent a confident per-residue confidence score (90 > pLDDT > 70), yellow segments represent a low confidence score (pLDDT > 50) and orange segments represent a very low confidence per-residue confidence score (pLDDT< 50).

**Fig. S2.**
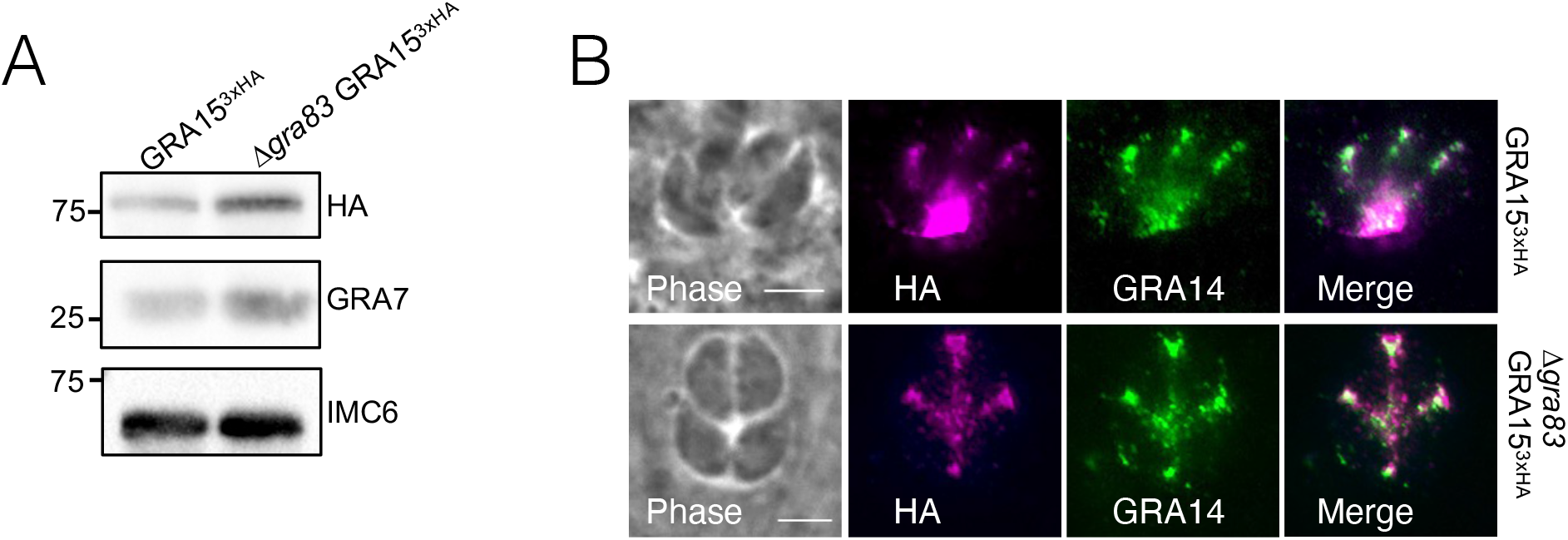
Deletion of *GRA83* does not impact GRA15 expression levels or localization. A) Western blot of whole cell lysates of GRA15^3xHA^ or Δ*gra83* GRA15^3xHA^ extracellular parasites probed with mouse anti-HA, mouse anti-GRA7, or rabbit anti-IMC6. B) IFA of GRA15^3xHA^ or Δ*gra83* GRA15^3xHA^ parasites showing that GRA15 localizes to the PV. Magenta: rabbit anti-HA; Green: mouse anti-GRA14.

**Fig. S3.**
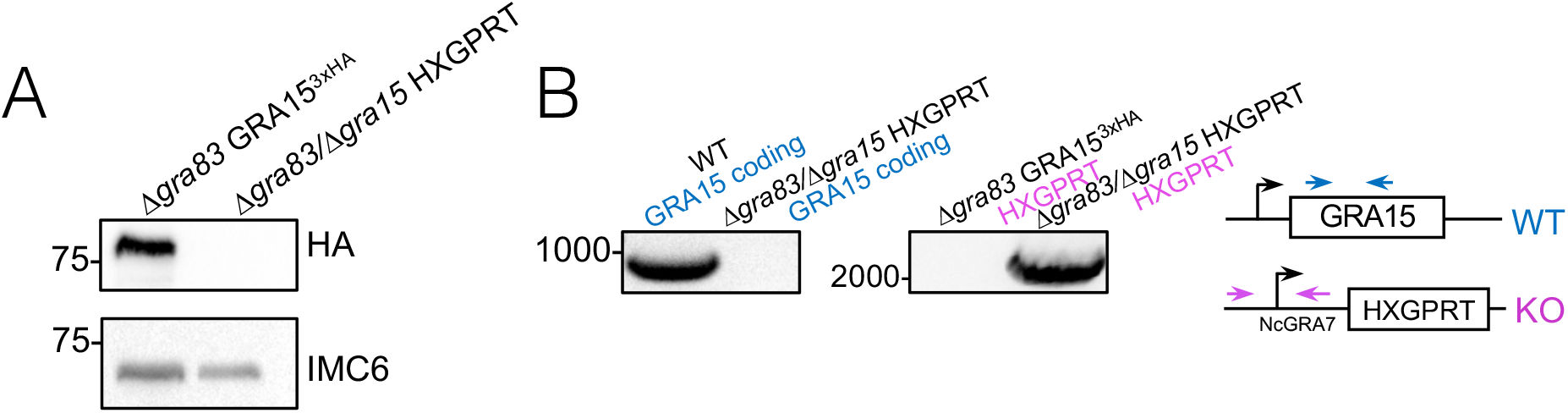
Verification of GRA15 disruption by western blot and PCR analyses. A) Western blot of the whole cell lysates of Δ*gra83* GRA15^3xHA^ or Δ*gra83/*Δ*gra15* strains shows the lack of HA signal in the Δ*gra83/*Δ*gra15* parasites. Probed with mouse anti-HA using rabbit anti-IMC6 as a load control. B) PCR verification of the removal of the *GRA15* coding region in the Δ*gra83/*Δ*gra15* strain and the confirmation of the *HXGPRT* recombination event. Diagram illustrates the positions of primers used to amplify the *GRA15* coding sequence (blue arrows) and knockout construct in the *GRA15* locus (magenta arrows).

**Figure S4.** List of hits identified by mass spectrometry of GRA83-TurboID whole cell lysates Proteins were ranked by spectral count. Two replicates of the untagged control (PruΔ*ku80*Δ*hxgprt)* and the GRA83^TurboID-3xHA^ are shown. Proteins not present in the PruΔ*ku80*Δ*hxgprt* control were ranked first, followed by proteins with the highest count in the GRA833^TurboID-3xHA^ samples. Gene IDs and descriptions are from ToxoDB.

**Table S1.**
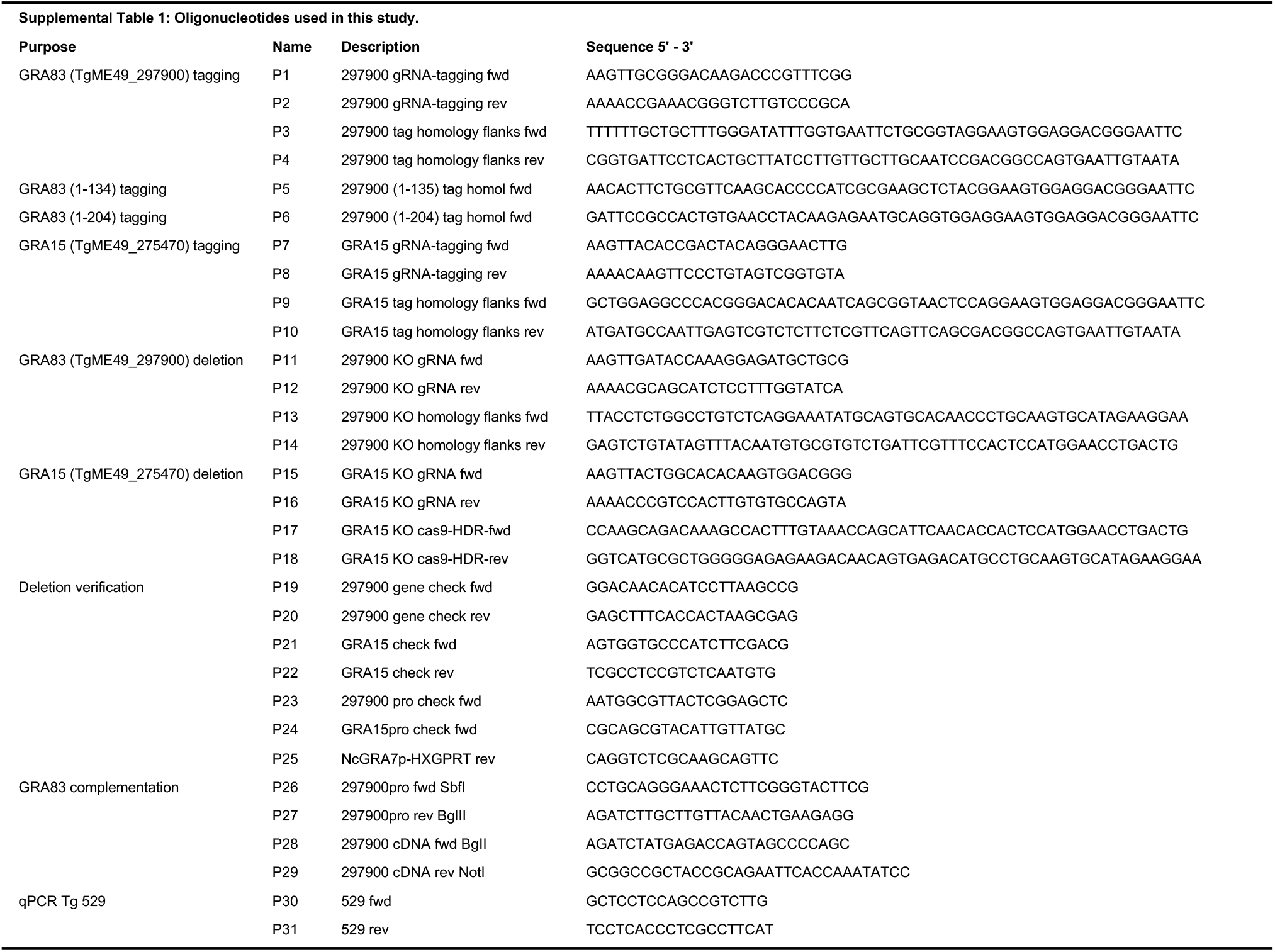
Oligonucleotide primers used in this study. All primer sequences are shown in the 5′-to-3′ orientation.

## References

1. Hill D, Dubey JP. 2002. Toxoplasma gondii: transmission, diagnosis and prevention. Clinical Microbiology and Infection 8:634–640.

2. Pappas G, Roussos N, Falagas ME. 2009. Toxoplasmosis snapshots: Global status of Toxoplasma gondii seroprevalence and implications for pregnancy and congenital toxoplasmosis. International Journal for Parasitology 39:1385–1394.

3. Smith NC, Goulart C, Hayward JA, Kupz A, Miller CM, van Dooren GG. 2021. Control of human toxoplasmosis. International Journal for Parasitology 51:95–121.

4. Porter SB, Sande MA. 1992. Toxoplasmosis of the Central Nervous System in the Acquired Immunodeficiency Syndrome. New England Journal of Medicine 327:1643–1648.

5. Wallace GR, Stanford MR. 2008. Immunity and Toxoplasma retinochoroiditis. Clin Exp Immunol 153:309–315.

6. Sullivan WJ, Jeffers V. 2012. Mechanisms of *Toxoplasma gondii* persistence and latency. FEMS Microbiol Rev 36:717–733.

7. Jones JL, Bonetti V, Holland GN, Press C, Sanislo SR, Khurana RN, Montoya JG. 2015. Ocular Toxoplasmosis in the United States: Recent and Remote Infections. Clinical Infectious Diseases 60:271–273.

8. Dubey JP, Lindsay DS, Speer CA. 1998. Structures of Toxoplasma gondii tachyzoites, bradyzoites, and sporozoites and biology and development of tissue cysts. Clin Microbiol Rev 11:267–299.

9. Dupont CD, Christian DA, Hunter CA. 2012. Immune response and immunopathology during toxoplasmosis. Semin Immunopathol 34:793–813.

10. Remington JS, Merigan TC. 1968. Interferon: Protection of Cells Infected with an Intracellular Protozoan (Toxoplasma gondii). Science 161:804–806.

11. Suzuki Y, Orellana MA, Schreiber RD, Remington JS. 1988. Interferon-$\gamma $: The Major Mediator of Resistance against Toxoplasma gondii. Science 240:516–518.

12. Suzuki Y, Conley FK, Remington JS. 1989. Importance of endogenous IFN-gamma for prevention of toxoplasmic encephalitis in mice. The Journal of Immunology 143:2045– 2050.

13. Pifer R, Yarovinsky F. 2011. Innate responses to Toxoplasma gondii in mice and humans. Trends Parasitol 27:388–393.

14. Yarovinsky F, Zhang D, Andersen JF, Bannenberg GL, Serhan CN, Hayden MS, Hieny S, Sutterwala FS, Flavell RA, Ghosh S, Sher A. 2005. TLR11 activation of dendritic cells by a protozoan profilin-like protein. Science 308:1626–1629.

15. Recognition of Profilin by Toll-like Receptor 12 Is Critical for Host Resistance to Toxoplasma gondii | Elsevier Enhanced Reader. https://reader.elsevier.com/reader/sd/pii/S1074761312005158?token=8B83A7354419BB8FCE59C61AE8AB3A2F455AACF3C66AFF7909480EABE62DC3E57FC8E4AE6598CE6F302AE3A72D443FF3&originRegion=us-east-1&originCreation=20230417185049. Retrieved 17 April 2023.

16. Andrade WA, Souza M do C, Ramos-Martinez E, Nagpal K, Dutra MS, Melo MB, Bartholomeu DC, Ghosh S, Golenbock DT, Gazzinelli RT. 2013. Combined Action of Nucleic Acid-Sensing Toll-like Receptors and TLR11/TLR12 Heterodimers Imparts Resistance to Toxoplasma gondii in Mice. Cell Host & Microbe 13:42–53.

17. Gazzinelli RT, Hieny S, Wynn TA, Wolf S, Sher A. 1993. Interleukin 12 is required for the T-lymphocyte-independent induction of interferon gamma by an intracellular parasite and induces resistance in T-cell-deficient hosts. Proc Natl Acad Sci U S A 90:6115–6119.

18. Hunter CA, Subauste CS, Van Cleave VH, Remington JS. 1994. Production of gamma interferon by natural killer cells from Toxoplasma gondii-infected SCID mice: regulation by interleukin-10, interleukin-12, and tumor necrosis factor alpha. Infection and Immunity 62:2818–2824.

19. Robben PM, Mordue DG, Truscott SM, Takeda K, Akira S, Sibley LD. 2004. Production of IL-12 by Macrophages Infected with Toxoplasma gondii Depends on the Parasite Genotype1. The Journal of Immunology 172:3686–3694.

20. Adams LB, Hibbs JB, Taintor RR, Krahenbuhl JL. 1990. Microbiostatic effect of murine-activated macrophages for Toxoplasma gondii. Role for synthesis of inorganic nitrogen oxides from L-arginine. The Journal of Immunology 144:2725–2729.

21. Pfefferkorn ER. 1984. Interferon gamma blocks the growth of Toxoplasma gondii in human fibroblasts by inducing the host cells to degrade tryptophan. Proceedings of the National Academy of Sciences 81:908–912.

22. Martens S, Parvanova I, Zerrahn J, Griffiths G, Schell G, Reichmann G, Howard JC. 2005. Disruption of Toxoplasma gondii Parasitophorous Vacuoles by the Mouse p47-Resistance GTPases. PLOS Pathogens 1:e24.

23. Hakimi M-A, Olias P, Sibley LD. 2017. Toxoplasma Effectors Targeting Host Signaling and Transcription. Clin Microbiol Rev 30:615–645.

24. Wang Y, Sangaré LO, Paredes-Santos TC, Saeij JPJ. 2020. Toxoplasma Mechanisms for Delivery of Proteins and Uptake of Nutrients Across the Host-Pathogen Interface. Annu Rev Microbiol 74:567–586.

25. Bougdour A, Durandau E, Brenier-Pinchart M-P, Ortet P, Barakat M, Kieffer S, Curt-Varesano A, Curt-Bertini R-L, Bastien O, Coute Y, Pelloux H, Hakimi M-A. 2013. Host Cell Subversion by Toxoplasma GRA16, an Exported Dense Granule Protein that Targets the Host Cell Nucleus and Alters Gene Expression. Cell Host & Microbe 13:489–500.

26. Panas MW, Boothroyd JC. 2020. Toxoplasma Uses GRA16 To Upregulate Host c-Myc. mSphere 5:e00402–20.

27. Rudzki EN, Ander SE, Coombs RS, Alrubaye HS, Cabo LF, Blank ML, Gutiérrez-Melo N, Dubey JP, Coyne CB, Boyle JP. 2021. Toxoplasma gondii *GRA28* Is Required for Placenta-Specific Induction of the Regulatory Chemokine CCL22 in Human and Mouse. mBio 12:e01591–21.

28. ten Hoeve AL, Braun L, Rodriguez ME, Olivera GC, Bougdour A, Belmudes L, Couté Y, Saeij JPJ, Hakimi M-A, Barragan A. 2022. The Toxoplasma effector GRA28 promotes parasite dissemination by inducing dendritic cell-like migratory properties in infected macrophages. Cell Host & Microbe 30:1570–1588.e7.

29. Braun L, Brenier-Pinchart M-P, Yogavel M, Curt-Varesano A, Curt-Bertini R-L, Hussain T, Kieffer-Jaquinod S, Coute Y, Pelloux H, Tardieux I, Sharma A, Belrhali H, Bougdour A, Hakimi M-A. 2013. A Toxoplasma dense granule protein, GRA24, modulates the early immune response to infection by promoting a direct and sustained host p38 MAPK activation. J Exp Med 210:2071–2086.

30. Rosowski EE, Lu D, Julien L, Rodda L, Gaiser RA, Jensen KDC, Saeij JPJ. 2011. Strain-specific activation of the NF-κB pathway by GRA15, a novel Toxoplasma gondii dense granule protein. J Exp Med 208:195–212.

31. Sangaré LO, Yang N, Konstantinou EK, Lu D, Mukhopadhyay D, Young LH, Saeij JPJ. 2019. Toxoplasma GRA15 Activates the NF-κB Pathway through Interactions with TNF Receptor-Associated Factors. mBio 10:e00808–19.

32. Mukhopadhyay D, Sangaré LO, Braun L, Hakimi M-A, Saeij JP. 2020. Toxoplasma GRA15 limits parasite growth in IFNγ-activated fibroblasts through TRAF ubiquitin ligases. EMBO J 39:e103758.

33. Nadipuram SM, Thind AC, Rayatpisheh S, Wohlschlegel JA, Bradley PJ. 2020. Proximity biotinylation reveals novel secreted dense granule proteins of Toxoplasma gondii bradyzoites. PLoS ONE 15:e0232552.

34. Barylyuk K, Koreny L, Ke H, Butterworth S, Crook OM, Lassadi I, Gupta V, Tromer E, Mourier T, Stevens TJ, Breckels LM, Pain A, Lilley KS, Waller RF. 2020. A Comprehensive Subcellular Atlas of the Toxoplasma Proteome via hyperLOPIT Provides Spatial Context for Protein Functions. Cell Host & Microbe 28:752–766.e9.

35. Harb OS, Roos DS. 2020. ToxoDB: Functional Genomics Resource for Toxoplasma and Related Organisms. Methods Mol Biol 2071:27–47.

36. Duvaud S, Gabella C, Lisacek F, Stockinger H, Ioannidis V, Durinx C. 2021. Expasy, the Swiss Bioinformatics Resource Portal, as designed by its users. Nucleic Acids Research 49:W216–W227.

37. Jumper J, Evans R, Pritzel A, Green T, Figurnov M, Ronneberger O, Tunyasuvunakool K, Bates R, Žídek A, Potapenko A, Bridgland A, Meyer C, Kohl SAA, Ballard AJ, Cowie A, Romera-Paredes B, Nikolov S, Jain R, Adler J, Back T, Petersen S, Reiman D, Clancy E, Zielinski M, Steinegger M, Pacholska M, Berghammer T, Bodenstein S, Silver D, Vinyals O, Senior AW, Kavukcuoglu K, Kohli P, Hassabis D. 2021. Highly accurate protein structure prediction with AlphaFold. 7873. Nature 596:583–589.

38. Varadi M, Anyango S, Deshpande M, Nair S, Natassia C, Yordanova G, Yuan D, Stroe O, Wood G, Laydon A, Žídek A, Green T, Tunyasuvunakool K, Petersen S, Jumper J, Clancy E, Green R, Vora A, Lutfi M, Figurnov M, Cowie A, Hobbs N, Kohli P, Kleywegt G, Birney E, Hassabis D, Velankar S. 2022. AlphaFold Protein Structure Database: massively expanding the structural coverage of protein-sequence space with high-accuracy models. Nucleic Acids Research 50:D439–D444.

39. Fox BA, Falla A, Rommereim LM, Tomita T, Gigley JP, Mercier C, Cesbron-Delauw M-F, Weiss LM, Bzik DJ. 2011. Type II Toxoplasma gondii KU80 Knockout Strains Enable Functional Analysis of Genes Required for Cyst Development and Latent Infection ▿. Eukaryot Cell 10:1193–1206.

40. Singh U, Brewer JL, Boothroyd JC. 2002. Genetic analysis of tachyzoite to bradyzoite differentiation mutants in Toxoplasma gondii reveals a hierarchy of gene induction. Molecular Microbiology 44:721–733.

41. Rome ME, Beck JR, Turetzky JM, Webster P, Bradley PJ. 2008. Intervacuolar Transport and Unique Topology of GRA14, a Novel Dense Granule Protein in Toxoplasma gondii. Infection and Immunity 76:4865–4875.

42. Pittman KJ, Aliota MT, Knoll LJ. 2014. Dual transcriptional profiling of mice and Toxoplasma gondii during acute and chronic infection. BMC Genomics 15:806.

43. Chern JH, Pasquarelli RR, Moon AS, Chen AL, Sha J, Wohlschlegel JA, Bradley PJ. 2021. A Novel Toxoplasma Inner Membrane Complex Suture-Associated Protein Regulates Suture Protein Targeting and Colocalizes with Membrane Trafficking Machinery. mBio 12:e02455–21.

44. Donald RG, Roos DS. 1995. Insertional mutagenesis and marker rescue in a protozoan parasite: cloning of the uracil phosphoribosyltransferase locus from Toxoplasma gondii. Proc Natl Acad Sci U S A 92:5749–5753.

45. Tissue tropism and parasite burden of Toxoplasma gondii RH strain in experimentally infected mice | Elsevier Enhanced Reader. https://reader.elsevier.com/reader/sd/pii/S1995764514600870?token=0BFDE4DB4B1BA0FB117E4608A0612E61ADEA737EB834FB05202479D9025485C4CBBD0AFD0976F92B00DDCCC6D64956DD&originRegion=us-east-1&originCreation=20230512041730. Retrieved 11 May 2023.

46. Lee YH, Kasper LH. 2004. Immune responses of different mouse strains after challenge with equivalent lethal doses of toxoplasma Gondii. 1. Parasite 11:89–97.

47. Butcher BA, Kim L, Johnson PF, Denkers EY. 2001. Toxoplasma gondii tachyzoites inhibit proinflammatory cytokine induction in infected macrophages by preventing nuclear translocation of the transcription factor NF-kappa B. J Immunol 167:2193–2201.

48. Krishnamurthy S, Maru P, Wang Y, Bitew MA, Mukhopadhyay D, Yamaryo-Botté Y, Paredes-Santos TC, Sangaré LO, Swale C, Botté CY, Saeij JPJ. 2023. CRISPR Screens Identify Toxoplasma Genes That Determine Parasite Fitness in Interferon Gamma-Stimulated Human Cells. mBio 0:e00060–23.

49. Nadipuram SM, Kim EW, Vashisht AA, Lin AH, Bell HN, Coppens I, Wohlschlegel JA, Bradley PJ. 2016. In Vivo Biotinylation of the Toxoplasma Parasitophorous Vacuole Reveals Novel Dense Granule Proteins Important for Parasite Growth and Pathogenesis. mBio 7:e00808–16.

50. Lecordier L, Moleon-Borodowsky I, Dubremetz J-F, Tourvieille B, Mercier C, Deslée D, Capron A, Cesbron-Delauw M-F. 1995. Characterization of a dense granule antigen of Toxoplasma gondii (GRA6) associated to the network of the parasitophorous vacuole. Molecular and Biochemical Parasitology 70:85–94.

51. Gay G, Braun L, Brenier-Pinchart M-P, Vollaire J, Josserand V, Bertini R-L, Varesano A, Touquet B, De Bock P-J, Coute Y, Tardieux I, Bougdour A, Hakimi M-A. 2016. Toxoplasma gondii TgIST co-opts host chromatin repressors dampening STAT1-dependent gene regulation and IFN-γ–mediated host defenses. Journal of Experimental Medicine 213:1779–1798.

52. Zheng X-N, Wang J-L, Elsheikha HM, Wang M, Zhang Z-W, Sun L-X, Wang X-C, Zhu X-Q, Li T-T. Functional Characterization of 15 Novel Dense Granule Proteins in Toxoplasma gondii Using the CRISPR-Cas9 System. Microbiol Spectr 11:e03078–22.

53. Levray YS, Bana B, Tarr SJ, McLaughlin EJ, Rossi-Smith P, Waltho A, Charlton GH, Chiozzi RZ, Straton CR, Thalassinos K, Osborne AR. 2023. Formation of ER-lumenal intermediates during export of Plasmodium proteins containing transmembrane-like hydrophobic sequences. PLOS Pathogens 19:e1011281.

54. Sidik SM, Huet D, Ganesan SM, Huynh M-H, Wang T, Nasamu AS, Thiru P, Saeij JPJ, Carruthers VB, Niles JC, Lourido S. 2016. A Genome-wide CRISPR Screen in Toxoplasma Identifies Essential Apicomplexan Genes. Cell 166:1423–1435.e12.

55. Jensen KDC, Wang Y, Wojno EDT, Shastri AJ, Hu K, Cornel L, Boedec E, Ong Y-C, Chien Y, Hunter CA, Boothroyd JC, Saeij JPJ. 2011. Toxoplasma polymorphic effectors determine macrophage polarization and intestinal inflammation. Cell Host Microbe 9:472– 483.

56. Ihara F, Fereig RM, Himori Y, Kameyama K, Umeda K, Tanaka S, Ikeda R, Yamamoto M, Nishikawa Y. 2020. Toxoplasma gondii Dense Granule Proteins 7, 14, and 15 Are Involved in Modification and Control of the Immune Response Mediated via NF-κB Pathway. Front Immunol 11:1709.

57. Wang P, Li S, Zhao Y, Zhang B, Li Y, Liu S, Du H, Cao L, Ou M, Ye X, Li P, Gao X, Wang P, Jing C, Shao F, Yang G, You F. 2019. The GRA15 protein from Toxoplasma gondii enhances host defense responses by activating the interferon stimulator STING. J Biol Chem 294:16494–16508.

58. Jensen KDC, Hu K, Whitmarsh RJ, Hassan MA, Julien L, Lu D, Chen L, Hunter CA, Saeij JPJ. 2013. Toxoplasma gondii rhoptry 16 kinase promotes host resistance to oral infection and intestinal inflammation only in the context of the dense granule protein GRA15. Infect Immun 81:2156–2167.

59. Fox BA, Guevara RB, Rommereim LM, Falla A, Bellini V, Pètre G, Rak C, Cantillana V, Dubremetz J-F, Cesbron-Delauw M-F, Taylor GA, Mercier C, Bzik DJ. 2019. *Toxoplasma gondii* Parasitophorous Vacuole Membrane-Associated Dense Granule Proteins Orchestrate Chronic Infection and GRA12 Underpins Resistance to Host Gamma Interferon. mBio 10:e00589–19, /mbio/10/4/mBio.00589-19.atom.

60. Gov L, Karimzadeh A, Ueno N, Lodoen MB. 2013. Human innate immunity to Toxoplasma gondii is mediated by host caspase-1 and ASC and parasite GRA15. mBio 4:e00255–13.

61. Braun L, Brenier-Pinchart M-P, Hammoudi P-M, Cannella D, Kieffer-Jaquinod S, Vollaire J, Josserand V, Touquet B, Couté Y, Tardieux I, Bougdour A, Hakimi M-A. 2019. The Toxoplasma effector TEEGR promotes parasite persistence by modulating NF-κB signalling via EZH2. Nat Microbiol 4:1208–1220.

62. Melo MB, Nguyen QP, Cordeiro C, Hassan MA, Yang N, McKell R, Rosowski EE, Julien L, Butty V, Dardé M-L, Ajzenberg D, Fitzgerald K, Young LH, Saeij JPJ. 2013. Transcriptional Analysis of Murine Macrophages Infected with Different Toxoplasma Strains Identifies Novel Regulation of Host Signaling Pathways. PLOS Pathogens 9:e1003779.

63. Herman RK, Molestina RE, Sinai AP, Howe DK. 2007. The apicomplexan pathogen Neospora caninum inhibits host cell apoptosis in the absence of discernible NF-kappa B activation. Infect Immun 75:4255–4262.

64. Mineo TWP, Chern JH, Thind AC, Mota CM, Nadipuram SM, Torres JA, Bradley PJ. 2022. Efficient Gene Knockout and Knockdown Systems in Neospora caninum Enable Rapid Discovery and Functional Assessment of Novel Proteins. mSphere 7:e00896–21.

65. Donald RG, Roos DS. 1993. Stable molecular transformation of Toxoplasma gondii: a selectable dihydrofolate reductase-thymidylate synthase marker based on drug-resistance mutations in malaria. Proceedings of the National Academy of Sciences 90:11703–11707.

66. Donald RGK, Carter D, Ullman B, Roos DS. 1996. Insertional Tagging, Cloning, and Expression of the Toxoplasma gondii Hypoxanthine-Xanthine-Guanine Phosphoribosyltransferase Gene: USE AS A SELECTABLE MARKER FOR STABLE TRANSFORMATION *. Journal of Biological Chemistry 271:14010–14019.

67. Viswanathan S, Williams ME, Bloss EB, Stasevich TJ, Speer CM, Nern A, Pfeiffer BD, Hooks BM, Li W-P, English BP, Tian T, Henry GL, Macklin JJ, Patel R, Gerfen CR, Zhuang X, Wang Y, Rubin GM, Looger LL. 2015. High-performance probes for light and electron microscopy. Nat Methods 12:568–576.

68. Sohn CS, Cheng TT, Drummond ML, Peng ED, Vermont SJ, Xia D, Cheng SJ, Wastling JM, Bradley PJ. 2011. Identification of Novel Proteins in Neospora caninum Using an Organelle Purification and Monoclonal Antibody Approach. PLoS One 6:e18383.

69. Choi CP, Moon AS, Back PS, Jami-Alahmadi Y, Vashisht AA, Wohlschlegel JA, Bradley PJ. 2019. A photoactivatable crosslinking system reveals protein interactions in the Toxoplasma gondii inner membrane complex. PLOS Biology 17:e3000475.

70. Straub KW, Cheng SJ, Sohn CS, Bradley PJ. 2009. Novel components of the Apicomplexan moving junction reveal conserved and coccidia-restricted elements. Cell Microbiol 11:590–603.

71. Sidik SM, Hackett CG, Tran F, Westwood NJ, Lourido S. 2014. Efficient Genome Engineering of Toxoplasma gondii Using CRISPR/Cas9. PLOS ONE 9:e100450.

72. Pfefferkorn ER, Borotz SE. 1994. Toxoplasma gondii: Characterization of a Mutant Resistant to 6-Thioxanthine. Experimental Parasitology 79:374–382.

73. Mayoral J, Di Cristina M, Carruthers VB, Weiss LM. 2020. Toxoplasma gondii: Bradyzoite Differentiation In Vitro and In Vivo, p. 269–282. In Tonkin, CJ (ed.), Toxoplasma gondii: Methods and Protocols. Springer US, New York, NY.

74. Mineo TWP, Benevides L, Silva NM, Silva JS. 2009. Myeloid differentiation factor 88 is required for resistance to *Neospora caninum* infection. Vet Res 40:32.

75. Mota CM, Lima-Junior D de S, Ferreira França FB, Aguillón Torres JD, Barros P da SC, Santiago FM, Silva JS, Mineo JR, Zamboni DS, Mineo TWP. 2020. Interplay Between Reactive Oxygen Species and the Inflammasome Are Crucial for Restriction of Neospora caninum Replication. Frontiers in Cellular and Infection Microbiology 10.

76. Xu T, Park SK, Venable JD, Wohlschlegel JA, Diedrich JK, Cociorva D, Lu B, Liao L, Hewel J, Han X, Wong CCL, Fonslow B, Delahunty C, Gao Y, Shah H, Yates JR. 2015. ProLuCID: An improved SEQUEST-like algorithm with enhanced sensitivity and specificity. J Proteomics 129:16–24.

77. Tabb DL, McDonald WH, Yates JR. 2002. DTASelect and Contrast: tools for assembling and comparing protein identifications from shotgun proteomics. J Proteome Res 1:21–26.

78. Cociorva D, L. Tabb D, Yates JR. 2006. Validation of Tandem Mass Spectrometry Database Search Results Using DTASelect. Current Protocols in Bioinformatics 16:13.4.1-13.4.14.

