## Supplemental Figure 4 for "The *Toxoplasma gondii* effector GRA83 modulates the host’s innate immune response to regulate parasite infection"

| Gene | Description | Known GRAs | control 1 | control 2 | GRA83-TurboID-1 | GRA83-TurboID-2 |
|---|---|---|---|---|---|---|
| TGME49_297900 | GRA83 (bait) | GRA83 | 0 | 0 | 290 | 244 |
| TGME49_264660 | SAG-related sequence SRS44 | SRS44/CST1 | 0 | 0 | 247 | 228 |
| TGME49_217680 | hypothetical protein | GRA57 | 0 | 0 | 215 | 236 |
| TGME49_232600 | phospholipase, patatin family protein | PLP1 | 0 | 0 | 187 | 223 |
| TGME49_239752 | hypothetical protein | #N/A | 0 | 0 | 202 | 186 |
| TGME49_310010 | rhoptry neck protein RON1 | #N/A | 0 | 0 | 157 | 222 |
| TGME49_231960 | putative omega secalin | GRA28 | 0 | 0 | 183 | 178 |
| TGME49_270320 | protein phosphatase 2C domain-containing protein | PPMC3 | 0 | 0 | 183 | 160 |
| TGME49_215910 | hypothetical protein | #N/A | 0 | 0 | 164 | 148 |
| TGME49_229010 | rhoptry neck protein RON4 | #N/A | 0 | 0 | 146 | 155 |
| TGME49_204480 | thioredoxin domain-containing protein | #N/A | 0 | 0 | 169 | 129 |
| TGME49_306060 | rhoptry neck protein RON8 | #N/A | 0 | 0 | 138 | 159 |
| TGME49_312420 | hypothetical protein | GRA38 | 0 | 0 | 158 | 128 |
| TGME49_275470 | dense granule protein GRA15 | GRA15 | 0 | 0 | 113 | 168 |
| TGME49_311210 | hypothetical protein | #N/A | 0 | 0 | 145 | 133 |
| TGME49_310780 | dense granule protein GRA4 | GRA04 | 0 | 0 | 135 | 142 |
| TGME49_270240 | MAG1 protein | MAG1 | 0 | 0 | 140 | 132 |
| TGME49_292020 | GCC2 and GCC3 domain-containing protein | #N/A | 0 | 0 | 147 | 118 |
| TGME49_247195 | microneme protein MIC15 | #N/A | 0 | 0 | 115 | 135 |
| TGME49_254470 | hypothetical protein | MYR1 | 0 | 0 | 125 | 115 |
| TGME49_297745 | hypothetical protein | #N/A | 0 | 0 | 104 | 122 |
| TGME49_266435 | hypothetical protein | #N/A | 0 | 0 | 97 | 115 |
| TGME49_228170 | inner membrane complex protein IMC2A | GRA44 | 0 | 0 | 99 | 92 |
| TGME49_288650 | dense granule protein GRA12 | GRA12 | 0 | 0 | 90 | 100 |
| TGME49_212300 | hypothetical protein | GRA32 | 0 | 0 | 96 | 88 |
| TGME49_289540 | hypothetical protein | SFP1 | 0 | 0 | 90 | 93 |
| TGME49_261850 | putative helicase | #N/A | 0 | 0 | 94 | 86 |
| TGME49_233870 | hypothetical protein | #N/A | 0 | 0 | 91 | 86 |
| TGME49_309760 | hypothetical protein | GRA55 | 0 | 0 | 81 | 93 |
| TGME49_310790 | hypothetical protein | CST9 | 0 | 0 | 74 | 88 |
| TGME49_279420 | hypothetical protein | #N/A | 0 | 0 | 74 | 84 |
| TGME49_211460 | hypothetical protein | MYR4 | 0 | 0 | 78 | 78 |
| TGME49_311230 | hypothetical protein | #N/A | 0 | 0 | 83 | 66 |
| TGME49_262730 | rhoptry protein ROP16 | #N/A | 0 | 0 | 64 | 85 |
| TGME49_211680 | protein disulfide isomerase | #N/A | 0 | 0 | 71 | 75 |
| TGME49_224460 | putative aminopeptidase n | #N/A | 0 | 0 | 74 | 67 |
| TGME49_201780 | microneme protein MIC2 | #N/A | 0 | 0 | 71 | 68 |
| TGME49_233460 | SAG-related sequence SRS29B | #N/A | 0 | 0 | 54 | 79 |
| TGME49_202780 | rhoptry kinase family protein ROP25 | ROP25 | 0 | 0 | 73 | 58 |
| TGME49_204340 | hypothetical protein | GRA54 (CST8) | 0 | 0 | 62 | 67 |
| TGME49_238040 | #N/A | #N/A | 0 | 0 | 49 | 76 |
| TGME49_229630 | eIF2 kinase IF2K-A (incomplete catalytic triad) | #N/A | 0 | 0 | 59 | 66 |
| TGME49_300100 | rhoptry neck protein RON2 | #N/A | 0 | 0 | 56 | 67 |
| TGME49_270250 | dense granule protein GRA1 | GRA01 | 0 | 0 | 81 | 42 |
| TGME49_297960 | #N/A | #N/A | 0 | 0 | 62 | 59 |
| TGME49_258870 | #N/A | CST7 | 0 | 0 | 56 | 65 |
| TGME49_245670 | pyruvate dehydrogenase complex subunit PDH-E1Alpha | #N/A | 0 | 0 | 52 | 64 |
| TGME49_226830 | DnaK family protein | #N/A | 0 | 0 | 55 | 60 |
| TGME49_221870 | hypothetical protein | #N/A | 0 | 0 | 51 | 61 |
| TGME49_314500 | subtilisin SUB2 | #N/A | 0 | 0 | 46 | 62 |
| TGME49_308970 | hypothetical protein | GRA12D | 0 | 0 | 55 | 53 |
| TGME49_208830 | hypothetical protein | GRA16 | 0 | 0 | 57 | 51 |
| TGME49_321640 | cell division protein CDC48AP | #N/A | 0 | 0 | 54 | 51 |
| TGME49_226072 | Ser/Thr phosphatase family protein | #N/A | 0 | 0 | 56 | 49 |
| TGME49_309930 | melibiase subfamily protein | GRA56 | 0 | 0 | 50 | 52 |
| TGME49_253370 | hypothetical protein | #N/A | 0 | 0 | 50 | 50 |
| TGME49_229480 | putative calcium binding protein precursor | #N/A | 0 | 0 | 43 | 55 |
| TGME49_251540 | dense granule protein GRA9 | GRA09 | 0 | 0 | 38 | 59 |
| TGME49_306890 | hypothetical protein | #N/A | 0 | 0 | 56 | 40 |
| TGME49_286680 | hypothetical protein | #N/A | 0 | 0 | 40 | 54 |
| TGME49_216060 | hypothetical protein | #N/A | 0 | 0 | 48 | 46 |
| TGME49_304990 | guanylate-binding protein, N-terminal domain-containing | #N/A | 0 | 0 | 40 | 48 |
| TGME49_203310 | dense granule protein GRA7 | GRA07 | 0 | 0 | 41 | 45 |
| TGME49_269950 | hypothetical protein | GRA61 | 0 | 0 | 42 | 43 |
| TGME49_309600 | hypothetical protein | GRA71 | 0 | 0 | 41 | 43 |
| TGME49_244380 | cactin | #N/A | 0 | 0 | 38 | 46 |
| TGME49_320080 | hypothetical protein | #N/A | 0 | 0 | 41 | 42 |
| TGME49_293590 | putative 3-oxoacyl-acyl-carrier protein synthase I/II | #N/A | 0 | 0 | 36 | 46 |
| TGME49_211260 | rhoptry kinase family protein ROP26 (incomplete catalyti | #N/A | 0 | 0 | 42 | 39 |
| TGME49_254490 | Sel1 repeat-containing protein | #N/A | 0 | 0 | 32 | 47 |
| TGME49_238170 | hypothetical protein | #N/A | 0 | 0 | 37 | 42 |
| TGME49_261080 | kringle domain-containing protein | #N/A | 0 | 0 | 38 | 40 |

| | | | | | | |
|---|---|---|---|---|---|---|
| TGME49_314020 | hypothetical protein | #N/A | 0 | 0 | 32 | 45 |
| TGME49_240090 | putative rhoptry kinase family protein ROP34 | ROP34/WNG2 | 0 | 0 | 33 | 43 |
| TGME49_210778 | hemimethylated DNA binding domain-containing protein | #N/A | 0 | 0 | 35 | 40 |
| TGME49_283850 | peptidyl-prolyl cis-trans isomerase | #N/A | 0 | 0 | 33 | 41 |
| TGME49_203600 | hypothetical protein | GRA50 (CST2) | 0 | 0 | 36 | 38 |
| TGME49_239740 | dense granule protein GRA14 | GRA14 | 0 | 0 | 36 | 37 |
| TGME49_226380 | hypothetical protein | GRA35 | 0 | 0 | 30 | 42 |
| TGME49_304955 | serine/threonine specific protein phosphatase | PPM11C | 0 | 0 | 30 | 41 |
| TGME49_244530 | hypothetical protein | GRA49 | 0 | 0 | 37 | 34 |
| TGME49_240060 | hypothetical protein | TgIST1 | 0 | 0 | 45 | 26 |
| TGME49_289380 | hypothetical protein | GRA39 | 0 | 0 | 39 | 29 |
| TGME49_204050 | subtilisin SUB1 | #N/A | 0 | 0 | 30 | 36 |
| TGME49_293180 | NADP-specific glutamate dehydrogenase | #N/A | 0 | 0 | 30 | 34 |
| TGME49_291960 | rhoptry kinase family protein ROP40 (incomplete catalyti | #N/A | 0 | 0 | 30 | 34 |
| TGME49_268950 | hypothetical protein | #N/A | 0 | 0 | 26 | 38 |
| TGME49_208740 | putative microneme protein | MCP3 | 0 | 0 | 38 | 26 |
| TGME49_233480 | SAG-related sequence SRS29C | #N/A | 0 | 0 | 28 | 35 |
| TGME49_263220 | rhoptry kinase family protein ROP21 | #N/A | 0 | 0 | 30 | 32 |
| TGME49_209950 | putative thioredoxin | #N/A | 0 | 0 | 31 | 30 |
| TGME49_280400 | hypothetical protein | #N/A | 0 | 0 | 29 | 31 |
| TGME49_249990 | hypothetical protein | GRA70 | 0 | 0 | 27 | 33 |
| TGME49_249270 | putative protein disulfide isomerase-related protein (pro | #N/A | 0 | 0 | 30 | 30 |
| TGME49_245490 | microneme protein MIC8 | #N/A | 0 | 0 | 24 | 36 |
| TGME49_308020 | SAG-related sequence SRS57 | #N/A | 0 | 0 | 32 | 27 |
| TGME49_279100 | hypothetical protein | MAF1 related pı | 0 | 0 | 20 | 37 |
| TGME49_318525 | hypothetical protein | #N/A | 0 | 0 | 29 | 27 |
| TGME49_297880 | dense granule protein DG32 | GRA23 | 0 | 0 | 28 | 28 |
| TGME49_277270 | NTPase II | NTPase II | 0 | 0 | 23 | 33 |
| TGME49_238100 | transmembrane protein | #N/A | 0 | 0 | 29 | 27 |
| TGME49_228130 | hypothetical protein | #N/A | 0 | 0 | 24 | 32 |
| TGME49_208560 | carrier superfamily protein | #N/A | 0 | 0 | 27 | 29 |
| TGME49_254880 | Alpha-galactosidase | #N/A | 0 | 0 | 21 | 34 |
| TGME49_249670 | cathepsin B | #N/A | 0 | 0 | 24 | 30 |
| TGME49_248990 | hypothetical protein | #N/A | 0 | 0 | 23 | 30 |
| TGME49_244120 | hypothetical protein | #N/A | 0 | 0 | 25 | 28 |
| TGME49_286630 | redoxin domain-containing protein | #N/A | 0 | 0 | 23 | 29 |
| TGME49_267740 | hypothetical protein | GRA48 | 0 | 0 | 24 | 28 |
| TGME49_230705 | hypothetical protein | GRA51 (CST3) | 0 | 0 | 22 | 30 |
| TGME49_220240 | hypothetical protein | GRA31 | 0 | 0 | 28 | 24 |
| TGME49_215220 | hypothetical protein | GRA22 | 0 | 0 | 25 | 27 |
| TGME49_288000 | hypothetical protein | #N/A | 0 | 0 | 23 | 27 |
| TGME49_272695 | hypothetical protein | #N/A | 0 | 0 | 24 | 26 |
| TGME49_226690 | hypothetical protein | #N/A | 0 | 0 | 26 | 24 |
| TGME49_242020 | mannosyl-oligosaccharide glucosidase | #N/A | 0 | 0 | 24 | 24 |
| TGME49_227810 | rhoptry kinase family protein ROP11 (incomplete catalyti | #N/A | 0 | 0 | 21 | 27 |
| TGME49_313080 | hypothetical protein | #N/A | 0 | 0 | 21 | 26 |
| TGME49_262060 | hypothetical protein | #N/A | 0 | 0 | 19 | 28 |
| TGME49_307860 | hypothetical protein | #N/A | 0 | 0 | 24 | 22 |
| TGME49_258950 | lectin family protein | #N/A | 0 | 0 | 22 | 24 |
| TGME49_255260 | apical membrane antigen AMA1 | #N/A | 0 | 0 | 25 | 21 |
| TGME49_202620 | hypothetical protein | GRA64 | 0 | 0 | 25 | 21 |
| TGME49_224060 | putative thioredoxin | #N/A | 0 | 0 | 20 | 25 |
| TGME49_203290 | hypothetical protein | GRA34 | 0 | 0 | 20 | 25 |
| TGME49_232000 | hypothetical protein | GRA30 | 0 | 0 | 18 | 26 |
| TGME49_205040 | PGAP1 family protein | #N/A | 0 | 0 | 20 | 24 |
| TGME49_273445 | sufB/sufD domain-containing protein | #N/A | 0 | 0 | 22 | 21 |
| TGME49_227948 | peptidase M16 inactive domain-containing protein | #N/A | 0 | 0 | 18 | 25 |
| TGME49_225200 | hypothetical protein | #N/A | 0 | 0 | 21 | 22 |
| TGME49_254720 | dense granule protein GRA8 | GRA08 | 0 | 0 | 23 | 19 |
| TGME49_319560 | microneme protein MIC3 | #N/A | 0 | 0 | 20 | 21 |
| TGME49_290700 | hypothetical protein | GRA25 | 0 | 0 | 22 | 19 |
| TGME49_227620 | dense granule protein GRA2 | GRA02 | 0 | 0 | 14 | 27 |
| TGME49_218240 | hypothetical protein | #N/A | 0 | 0 | 28 | 13 |
| TGME49_201130 | rhoptry kinase family protein ROP33 | ROP33 | 0 | 0 | 23 | 18 |
| TGME49_299780 | hypothetical protein | #N/A | 0 | 0 | 19 | 21 |
| TGME49_269690 | hypothetical protein | GRA29 | 0 | 0 | 21 | 19 |
| TGME49_253900 | parasite porphobilinogen synthase PBGS | #N/A | 0 | 0 | 14 | 26 |
| TGME49_237015 | hypothetical protein | GRA43 | 0 | 0 | 11 | 29 |
| TGME49_232340 | protein phosphatase 2C domain-containing protein | #N/A | 0 | 0 | 19 | 21 |
| TGME49_214080 | toxofilin | #N/A | 0 | 0 | 21 | 19 |
| TGME49_291890 | microneme protein MIC1 | #N/A | 0 | 0 | 22 | 17 |
| TGME49_258770 | putative polypeptide n-acetylgalactosaminyltransferase 1 | #N/A | 0 | 0 | 19 | 20 |
| TGME49_237230 | hypothetical protein | MYR3 | 0 | 0 | 21 | 18 |

| | | | | | | |
|---|---|---|---|---|---|---|
| TGME49_237180 | hypothetical protein | #N/A | 0 | 0 | 20 | 19 |
| TGME49_316250 | hypothetical protein | GRA45 | 0 | 0 | 21 | 17 |
| TGME49_290160 | putative sortilin | #N/A | 0 | 0 | 16 | 22 |
| TGME49_263560 | hypothetical protein | #N/A | 0 | 0 | 17 | 21 |
| TGME49_239010 | hypothetical protein | TEEGR (HCE) | 0 | 0 | 20 | 18 |
| TGME49_205740 | hypothetical protein | #N/A | 0 | 0 | 17 | 21 |
| TGME49_258390 | putative DnaJ protein | #N/A | 0 | 0 | 18 | 19 |
| TGME49_253400 | hypothetical protein | #N/A | 0 | 0 | 18 | 19 |
| TGME49_319340 | hypothetical protein | GRA52 (CST5) | 0 | 0 | 17 | 19 |
| TGME49_308810 | #N/A | #N/A | 0 | 0 | 16 | 20 |
| TGME49_269885 | #N/A | #N/A | 0 | 0 | 16 | 20 |
| TGME49_258458 | hypothetical protein | #N/A | 0 | 0 | 14 | 22 |
| TGME49_300060 | signal peptidase subunit protein | #N/A | 0 | 0 | 18 | 17 |
| TGME49_266090 | hypothetical protein | #N/A | 0 | 0 | 17 | 18 |
| TGME49_228630 | hypothetical protein | #N/A | 0 | 0 | 15 | 20 |
| TGME49_262920 | trypsin domain-containing protein | #N/A | 0 | 0 | 16 | 18 |
| TGME49_262400 | lipase | #N/A | 0 | 0 | 16 | 18 |
| TGME49_247440 | hypothetical protein | GRA33 | 0 | 0 | 16 | 18 |
| TGME49_237500 | protein phosphatase 2C domain-containing protein | PPM3A | 0 | 0 | 18 | 16 |
| TGME49_208730 | putative microneme protein | MCP4 | 0 | 0 | 22 | 12 |
| TGME49_203010 | aurora kinase | #N/A | 0 | 0 | 16 | 18 |
| TGME49_286620 | S1 RNA binding domain-containing protein | #N/A | 0 | 0 | 10 | 23 |
| TGME49_271760 | seryl-tRNA synthetase (SerRS2) | #N/A | 0 | 0 | 15 | 18 |
| TGME49_240600 | putative chaperonin cpn60 | #N/A | 0 | 0 | 15 | 18 |
| TGME49_240430 | glyoxalase family protein | #N/A | 0 | 0 | 17 | 16 |
| TGME49_281980 | phosphatidate cytidylyltransferase | #N/A | 0 | 0 | 18 | 14 |
| TGME49_200360 | hypothetical protein | NcGRA47? | 0 | 0 | 16 | 16 |
| TGME49_314850 | hypothetical protein | #N/A | 0 | 0 | 14 | 17 |
| TGME49_297650 | Ser/Thr phosphatase family protein | #N/A | 0 | 0 | 20 | 11 |
| TGME49_247220 | nudix -type motif 9 isoform a family protein | #N/A | 0 | 0 | 17 | 14 |
| TGME49_246550 | aspartyl protease ASP3 | #N/A | 0 | 0 | 13 | 18 |
| TGME49_285180 | hypothetical protein | #N/A | 0 | 0 | 14 | 16 |
| TGME49_220890 | hypothetical protein | GRA12A | 0 | 0 | 16 | 14 |
| TGME49_206510 | toxolysin TLN4 | #N/A | 0 | 0 | 13 | 17 |
| TGME49_267730 | hypothetical protein | #N/A | 0 | 0 | 12 | 17 |
| TGME49_261750 | rhoptry neck protein RON10 | #N/A | 0 | 0 | 12 | 17 |
| TGME49_241240 | hypothetical protein | GRA21 | 0 | 0 | 18 | 11 |
| TGME49_204130 | perforin-like protein PLP1 | #N/A | 0 | 0 | 12 | 17 |
| TGME49_254000 | hypothetical protein | GRA47 | 0 | 0 | 11 | 17 |
| TGME49_253890 | peptidase M16 inactive domain-containing protein | #N/A | 0 | 0 | 10 | 18 |
| TGME49_230180 | hypothetical protein | GRA24 | 0 | 0 | 15 | 13 |
| TGME49_229490 | tetratricopeptide repeat-containing protein | #N/A | 0 | 0 | 9 | 19 |
| TGME49_260520 | hypothetical protein | GRA53 (CST6) | 0 | 0 | 14 | 13 |
| TGME49_258462 | hypothetical protein | TgPI-2 | 0 | 0 | 14 | 13 |
| TGME49_315320 | SAG-related sequence SRS52A | #N/A | 0 | 0 | 13 | 13 |
| TGME49_250955 | KRUF family protein | #N/A | 0 | 0 | 15 | 11 |
| TGME49_233450 | SAG-related sequence SRS29A | #N/A | 0 | 0 | 9 | 17 |
| TGME49_217740 | 3-ketoacyl-(acyl-carrier-protein) reductase | #N/A | 0 | 0 | 11 | 15 |
| TGME49_201180 | #N/A | MSF | 0 | 0 | 11 | 15 |
| TGME49_273960 | chaperonin GroS protein | #N/A | 0 | 0 | 13 | 12 |
| TGME49_266410 | hypothetical protein | #N/A | 0 | 0 | 13 | 12 |
| TGME49_258230 | rhoptry kinase family protein ROP20 | #N/A | 0 | 0 | 10 | 15 |
| TGME49_228360 | putative peptidyl-prolyl isomerase FKBP12 | #N/A | 0 | 0 | 10 | 15 |
| TGME49_222100 | hypothetical protein | #N/A | 0 | 0 | 13 | 12 |
| TGME49_312110 | apicoplast-associated thioredoxin family protein Atrx1 | #N/A | 0 | 0 | 13 | 11 |
| TGME49_275690 | putative ClpB | #N/A | 0 | 0 | 8 | 16 |
| TGME49_233220 | hypothetical protein | #N/A | 0 | 0 | 14 | 10 |
| TGME49_231130 | hypothetical protein | #N/A | 0 | 0 | 11 | 13 |
| TGME49_227280 | dense granule protein GRA3 | GRA03 | 0 | 0 | 11 | 13 |
| TGME49_308093 | rhoptry kinase family protein (incomplete catalytic triad) | ROP32 | 0 | 0 | 13 | 10 |
| TGME49_249820 | ATP-binding cassette sub-family B member 5 | #N/A | 0 | 0 | 14 | 9 |
| TGME49_248140 | hypothetical protein | #N/A | 0 | 0 | 8 | 15 |
| TGME49_246220 | hypothetical protein | #N/A | 0 | 0 | 11 | 12 |
| TGME49_219130 | NADPH-glutathione reductase | #N/A | 0 | 0 | 8 | 15 |
| TGME49_289000 | hypothetical protein | #N/A | 0 | 0 | 15 | 7 |
| TGME49_280380 | poly(ADP-ribose) glycohydrolase | #N/A | 0 | 0 | 7 | 15 |
| TGME49_262880 | hypothetical protein | #N/A | 0 | 0 | 10 | 12 |
| TGME49_222170 | dense-granule antigen DG32 | GRA17 | 0 | 0 | 12 | 10 |
| TGME49_211860 | hypothetical protein | #N/A | 0 | 0 | 13 | 9 |
| TGME49_320490 | N-acyl-phosphatidylethanolamine-hydrolyzing phospholip | GRA66 | 0 | 0 | 10 | 11 |
| TGME49_289360 | hypothetical protein | #N/A | 0 | 0 | 9 | 12 |
| TGME49_294220 | hypothetical protein | #N/A | 0 | 0 | 11 | 9 |
| TGME49_282055 | protein phosphatase PP2C-hn | #N/A | 0 | 0 | 10 | 10 |

| | | | | | | |
|---|---|---|---|---|---|---|
| TGME49_208030 | microneme protein MIC4 | #N/A | 0 | 0 | 10 | 10 |
| TGME49_305270 | hypothetical protein | #N/A | 0 | 0 | 8 | 11 |
| TGME49_300220 | hypothetical protein | #N/A | 0 | 0 | 11 | 8 |
| TGME49_272660 | hypothetical protein | #N/A | 0 | 0 | 10 | 9 |
| TGME49_215360 | hypothetical protein | GRA62 | 0 | 0 | 7 | 12 |
| TGME49_313640 | hypothetical protein | #N/A | 0 | 0 | 7 | 11 |
| TGME49_313270 | hypothetical protein | #N/A | 0 | 0 | 7 | 11 |
| TGME49_294360 | putative ubiquitin specific protease 39 isoform 2 | #N/A | 0 | 0 | 8 | 10 |
| TGME49_288840 | hypothetical protein | GRA18 | 0 | 0 | 8 | 10 |
| TGME49_280740 | signal peptidase | #N/A | 0 | 0 | 9 | 9 |
| TGME49_280370 | hypothetical protein | #N/A | 0 | 0 | 7 | 11 |
| TGME49_264200 | hypothetical protein | #N/A | 0 | 0 | 7 | 11 |
| TGME49_261780 | microneme protein MIC7 | #N/A | 0 | 0 | 7 | 11 |
| TGME49_254110 | tryptophanyl-tRNA synthetase (TrpRS1) | #N/A | 0 | 0 | 9 | 9 |
| TGME49_232020 | hypothetical protein | #N/A | 0 | 0 | 8 | 10 |
| TGME49_225850 | peptidase, M28 family protein | #N/A | 0 | 0 | 8 | 10 |
| TGME49_214940 | MIC2-associated protein M2AP | #N/A | 0 | 0 | 10 | 8 |
| TGME49_312200 | serine/threonine protein phosphatase | #N/A | 0 | 0 | 7 | 10 |
| TGME49_209060 | thrombospondin type 1 domain-containing protein | #N/A | 0 | 0 | 8 | 9 |
| TGME49_297710 | hypothetical protein | #N/A | 0 | 0 | 9 | 7 |
| TGME49_278770 | hypothetical protein | #N/A | 0 | 0 | 11 | 5 |
| TGME49_235340 | hypothetical protein | #N/A | 0 | 0 | 5 | 11 |
| TGME49_223420 | DnaJ domain-containing protein | #N/A | 0 | 0 | 6 | 10 |
| TGME49_205240 | cleft lip and palate transmembrane protein 1 (clptm1) pr | #N/A | 0 | 0 | 7 | 9 |
| TGME49_290730 | CS domain-containing protein | #N/A | 0 | 0 | 5 | 10 |
| TGME49_282170 | hypothetical protein | #N/A | 0 | 0 | 7 | 8 |
| TGME49_275440 | dense granule protein GRA6 | GRA06 | 0 | 0 | 6 | 9 |
| TGME49_233810 | Sel1 repeat-containing protein | #N/A | 0 | 0 | 8 | 7 |
| TGME49_230510 | hypothetical protein | #N/A | 0 | 0 | 9 | 6 |
| TGME49_225150 | hypothetical protein | #N/A | 0 | 0 | 7 | 8 |
| TGME49_219810 | hypothetical protein | GRA40 | 0 | 0 | 7 | 8 |
| TGME49_205250 | rhoptry protein ROP18 | #N/A | 0 | 0 | 6 | 9 |
| TGME49_204160 | GYF domain-containing protein | #N/A | 0 | 0 | 7 | 8 |
| TGME49_308000 | Gpi16 subunit, GPI transamidase component protein | #N/A | 0 | 0 | 5 | 9 |
| TGME49_300260 | threonyl-tRNA synthetase family protein | #N/A | 0 | 0 | 7 | 7 |
| TGME49_295090 | hypothetical protein | #N/A | 0 | 0 | 7 | 7 |
| TGME49_291180 | hypothetical protein | #N/A | 0 | 0 | 3 | 11 |
| TGME49_272290 | pyruvate dehydrogenase complex subunit PD-HE1Beta | #N/A | 0 | 0 | 5 | 9 |
| TGME49_262560 | hypothetical protein | #N/A | 0 | 0 | 6 | 8 |
| TGME49_258360 | hypothetical protein | #N/A | 0 | 0 | 7 | 7 |
| TGME49_232730 | acyl-CoA:diacylglycerol acyltransferase 1-related enzyme | #N/A | 0 | 0 | 5 | 9 |
| TGME49_226400 | lipoic acid synthase LIPA | #N/A | 0 | 0 | 4 | 10 |
| TGME49_224850 | putative polyadenylate binding protein | #N/A | 0 | 0 | 7 | 7 |
| TGME49_220330 | hypothetical protein | #N/A | 0 | 0 | 6 | 8 |
| TGME49_211440 | hypothetical protein | #N/A | 0 | 0 | 7 | 7 |
| TGME49_203970 | dolichyl-diphosphooligosaccharide--protein glycosyltransf | #N/A | 0 | 0 | 4 | 10 |
| TGME49_291630 | hypothetical protein | #N/A | 0 | 0 | 4 | 9 |
| TGME49_263270 | glycerophosphodiester phosphodiesterase family protein | #N/A | 0 | 0 | 5 | 8 |
| TGME49_256840 | hypothetical protein | #N/A | 0 | 0 | 7 | 6 |
| TGME49_251500 | putative eukaryotic initiation factor-3, subunit 3 | #N/A | 0 | 0 | 3 | 10 |
| TGME49_249340 | hypothetical protein | #N/A | 0 | 0 | 6 | 7 |
| TGME49_247400 | hypothetical protein | #N/A | 0 | 0 | 6 | 7 |
| TGME49_242230 | #N/A | #N/A | 0 | 0 | 4 | 9 |
| TGME49_242118 | #N/A | #N/A | 0 | 0 | 4 | 9 |
| TGME49_232050 | DnaJ domain-containing protein | #N/A | 0 | 0 | 5 | 8 |
| TGME49_231370 | phospholipase, patatin family protein | #N/A | 0 | 0 | 7 | 6 |
| TGME49_225160 | hypothetical protein | #N/A | 0 | 0 | 6 | 7 |
| TGME49_212140 | hypothetical protein | #N/A | 0 | 0 | 3 | 10 |
| TGME49_211290 | rhoptry protein ROP15 | #N/A | 0 | 0 | 8 | 5 |
| TGME49_206610 | pyruvate dehydrogenase complex subunit PDH-E2 | #N/A | 0 | 0 | 8 | 5 |
| TGME49_202940 | hypothetical protein | #N/A | 0 | 0 | 8 | 5 |
| TGME49_202610 | protein phosphatase 2C domain-containing protein | PPM3D | 0 | 0 | 7 | 6 |
| TGME49_323100 | #N/A | #N/A | 0 | 0 | 3 | 9 |
| TGME49_268690 | hypothetical protein | #N/A | 0 | 0 | 3 | 9 |
| TGME49_267590 | hypothetical protein | #N/A | 0 | 0 | 4 | 8 |
| TGME49_253030 | glycosyl hydrolase, family 31 protein | #N/A | 0 | 0 | 5 | 7 |
| TGME49_247000 | tetratricopeptide repeat-containing protein | #N/A | 0 | 0 | 5 | 7 |
| TGME49_236010 | prenylcysteine oxidase | #N/A | 0 | 0 | 5 | 7 |
| TGME49_222370 | SAG-related sequence SRS13 | SRS13 | 0 | 0 | 2 | 10 |
| TGME49_217400 | hypothetical protein | #N/A | 0 | 0 | 6 | 6 |
| TGME49_216720 | hypothetical protein | #N/A | 0 | 0 | 4 | 8 |
| TGME49_208450 | protease inhibitor PI2 | TgPI-1 | 0 | 0 | 8 | 4 |
| TGME49_205360 | hypothetical protein | #N/A | 0 | 0 | 4 | 8 |

| | | | | | | |
|---|---|---|---|---|---|---|
| TGME49_313690 | Sel1 repeat-containing protein | #N/A | 0 | 0 | 6 | 5 |
| TGME49_311690 | UBA/TS-N domain-containing protein | #N/A | 0 | 0 | 5 | 6 |
| TGME49_310430 | #N/A | #N/A | 0 | 0 | 3 | 8 |
| TGME49_308840 | SAG-related sequence SRS51 | #N/A | 0 | 0 | 5 | 6 |
| TGME49_294790 | hypothetical protein | #N/A | 0 | 0 | 7 | 4 |
| TGME49_279550 | hypothetical protein | #N/A | 0 | 0 | 6 | 5 |
| TGME49_272420 | putative phosphatidylcholine-sterol O-acyltransferase | LCAT | 0 | 0 | 7 | 4 |
| TGME49_247350 | thioredoxin domain-containing protein | #N/A | 0 | 0 | 5 | 6 |
| TGME49_230960 | putative splicing factor 3b, subunit 3 | #N/A | 0 | 0 | 2 | 9 |
| TGME49_204530 | microneme protein MIC11 | #N/A | 0 | 0 | 4 | 7 |
| TGME49_202572 | ribophorin i protein | #N/A | 0 | 0 | 4 | 7 |
| TGME49_201840 | aspartyl protease ASP1 | #N/A | 0 | 0 | 2 | 9 |
| TGME49_323110 | #N/A | #N/A | 0 | 0 | 5 | 5 |
| TGME49_319350 | SAG-related sequence SRS17B | #N/A | 0 | 0 | 8 | 2 |
| TGME49_297940 | single-strand binding protein | #N/A | 0 | 0 | 4 | 6 |
| TGME49_272460 | hypothetical protein | GRA72 | 0 | 0 | 5 | 5 |
| TGME49_254690 | phospholipase/carboxylesterase | #N/A | 0 | 0 | 5 | 5 |
| TGME49_246130 | serpin (serine proteinase inhibitor) superfamily protein | TgPI-1 | 0 | 0 | 5 | 5 |
| TGME49_227150 | putative glutaredoxin | #N/A | 0 | 0 | 4 | 6 |
| TGME49_224350 | #N/A | #N/A | 0 | 0 | 3 | 7 |
| TGME49_223940 | GAP45 protein | #N/A | 0 | 0 | 4 | 6 |
| TGME49_213880 | hypothetical protein | #N/A | 0 | 0 | 5 | 5 |
| TGME49_213067 | hypothetical protein | GRA36 | 0 | 0 | 5 | 5 |
| TGME49_208360 | hypothetical protein | #N/A | 0 | 0 | 4 | 6 |
| TGME49_204310 | hypothetical protein | #N/A | 0 | 0 | 4 | 6 |
| TGME49_201520 | protein phosphatase 2C domain-containing protein | #N/A | 0 | 0 | 5 | 5 |
| TGME49_306660 | RNA pseudouridine synthase superfamily protein | #N/A | 0 | 0 | 4 | 5 |
| TGME49_304740 | rhoptry kinase family protein ROP35 | ROP35/WNG1 | 0 | 0 | 3 | 6 |
| TGME49_231180 | hypothetical protein | #N/A | 0 | 0 | 4 | 5 |
| TGME49_216770 | hypothetical protein | #N/A | 0 | 0 | 4 | 5 |
| TGME49_211890 | RAP domain-containing protein | #N/A | 0 | 0 | 2 | 7 |
| TGME49_315910 | hypothetical protein | #N/A | 0 | 0 | 5 | 3 |
| TGME49_289580 | strictosidine synthase subfamily protein | #N/A | 0 | 0 | 3 | 5 |
| TGME49_277720 | GDA1/CD39 (nucleoside phosphatase) family protein | #N/A | 0 | 0 | 3 | 5 |
| TGME49_273730 | phospholipase, patatin family protein | #N/A | 0 | 0 | 3 | 5 |
| TGME49_254580 | UDP-galactose transporter family protein | #N/A | 0 | 0 | 3 | 5 |
| TGME49_251930 | enoyl-acyl carrier reductase ENR | #N/A | 0 | 0 | 4 | 4 |
| TGME49_246580 | hypothetical protein | #N/A | 0 | 0 | 4 | 4 |
| TGME49_244840 | zinc knuckle domain-containing protein | #N/A | 0 | 0 | 4 | 4 |
| TGME49_226960 | phosphofructokinase PFKII | #N/A | 0 | 0 | 4 | 4 |
| TGME49_218840 | mutS domain protein | #N/A | 0 | 0 | 3 | 5 |
| TGME49_217692 | hypothetical protein | #N/A | 0 | 0 | 2 | 6 |
| TGME49_215160 | hypothetical protein | #N/A | 0 | 0 | 3 | 5 |
| TGME49_214180 | ENTH domain-containing protein | #N/A | 0 | 0 | 4 | 4 |
| TGME49_214090 | signal peptidase | #N/A | 0 | 0 | 3 | 5 |
| TGME49_208040 | aldo-keto reductase | #N/A | 0 | 0 | 4 | 4 |
| TGME49_207120 | Sad1/UNC family protein | #N/A | 0 | 0 | 3 | 5 |
| TGME49_264970 | hypothetical protein | #N/A | 0 | 0 | 2 | 5 |
| TGME49_258826 | hypothetical protein | #N/A | 0 | 0 | 0 | 7 |
| TGME49_253330 | Rhoptry kinase family protein | BPK1 | 0 | 0 | 3 | 4 |
| TGME49_247680 | hypothetical protein | #N/A | 0 | 0 | 5 | 2 |
| TGME49_244180 | microneme-like protein | #N/A | 0 | 0 | 4 | 3 |
| TGME49_240280 | S1/P1nuclease | #N/A | 0 | 0 | 4 | 3 |
| TGME49_234950 | protein kinase (incomplete catalytic triad) | ROP48 | 0 | 0 | 4 | 3 |
| TGME49_231630 | alveolin domain containing intermediate filament IMC4 | #N/A | 0 | 0 | 4 | 3 |
| TGME49_221210 | cyclophilin | Cyclophilin | 0 | 0 | 4 | 3 |
| TGME49_215690 | hypothetical protein | #N/A | 0 | 0 | 4 | 3 |
| TGME49_213280 | SAG-related sequence SRS25 | #N/A | 0 | 0 | 3 | 4 |
| TGME49_297320 | hypothetical protein | #N/A | 0 | 0 | 4 | 2 |
| TGME49_271050 | SAG-related sequence SRS34A | #N/A | 0 | 0 | 0 | 6 |
| TGME49_269340 | hypothetical protein | #N/A | 0 | 0 | 0 | 6 |
| TGME49_266320 | hypothetical protein | #N/A | 0 | 0 | 3 | 3 |
| TGME49_250880 | kinase, pfkB family protein | #N/A | 0 | 0 | 0 | 6 |
| TGME49_247380 | hypothetical protein | #N/A | 0 | 0 | 0 | 6 |
| TGME49_321690 | hypothetical protein | #N/A | 0 | 0 | 3 | 2 |
| TGME49_313230 | eukaryotic initiation factor-2, alpha subunit | #N/A | 0 | 0 | 2 | 3 |
| TGME49_294250 | WD domain, G-beta repeat-containing protein | #N/A | 0 | 0 | 2 | 3 |
| TGME49_289150 | hypothetical protein | #N/A | 0 | 0 | 2 | 3 |
| TGME49_285870 | SAG-related sequence SRS20A | #N/A | 0 | 0 | 2 | 3 |
| TGME49_283780 | glucose-6-phosphate isomerase GPI | #N/A | 0 | 0 | 3 | 2 |
| TGME49_268790 | hypothetical protein | GRA58 | 0 | 0 | 2 | 3 |
| TGME49_260600 | Pumilio-family RNA binding repeat-containing protein | #N/A | 0 | 0 | 5 | 0 |
| TGME49_258730 | hypothetical protein | #N/A | 0 | 0 | 3 | 2 |

| | | | | | | |
|---|---|---|---|---|---|---|
| TGME49_255330 | hypothetical protein | #N/A | 0 | 0 | 5 | 0 |
| TGME49_245610 | hypothetical protein | #N/A | 0 | 0 | 2 | 3 |
| TGME49_243730 | rhoptry protein ROP9 | #N/A | 0 | 0 | 0 | 5 |
| TGME49_240970 | hypothetical protein | #N/A | 0 | 0 | 2 | 3 |
| TGME49_239600 | rhoptry kinase family protein ROP23 (incomplete catalyti | #N/A | 0 | 0 | 2 | 3 |
| TGME49_220270 | alveolin domain containing intermediate filament IMC6 | #N/A | 0 | 0 | 0 | 5 |
| TGME49_219250 | acetyltransferase, GNAT family protein | #N/A | 0 | 0 | 5 | 0 |
| TGME49_218540 | putative peptidase S15 | #N/A | 0 | 0 | 2 | 3 |
| TGME49_217530 | hypothetical protein | GRA63 | 0 | 0 | 0 | 5 |
| TGME49_213660 | zinc finger (CCCH type) motif-containing protein | #N/A | 0 | 0 | 3 | 2 |
| TGME49_315885 | putative glycosyltransferase | #N/A | 0 | 0 | 2 | 2 |
| TGME49_310150 | AMP-binding enzyme domain-containing protein | #N/A | 0 | 0 | 2 | 2 |
| TGME49_295990 | putative ubiquitin conjugating enzyme E2 | #N/A | 0 | 0 | 0 | 4 |
| TGME49_293570 | putative translocation protein sec62 | #N/A | 0 | 0 | 0 | 4 |
| TGME49_292920 | putative heat shock protein 75 | #N/A | 0 | 0 | 0 | 4 |
| TGME49_289650 | PEP-carboxykinase I | #N/A | 0 | 0 | 0 | 4 |
| TGME49_287270 | hypothetical protein | #N/A | 0 | 0 | 2 | 2 |
| TGME49_275460 | hypothetical protein | #N/A | 0 | 0 | 0 | 4 |
| TGME49_268200 | RNA recognition motif-containing protein | #N/A | 0 | 0 | 0 | 4 |
| TGME49_258590 | hypothetical protein | #N/A | 0 | 0 | 2 | 2 |
| TGME49_256990 | glycyl-tRNA synthetase | #N/A | 0 | 0 | 2 | 2 |
| TGME49_250010 | Sad1 / UNC family protein | #N/A | 0 | 0 | 0 | 4 |
| TGME49_249850 | GAP40 protein | #N/A | 0 | 0 | 0 | 4 |
| TGME49_219730 | hypothetical protein | #N/A | 0 | 0 | 0 | 4 |
| TGME49_210370 | hypothetical protein | #N/A | 0 | 0 | 2 | 2 |
| TGME49_307570 | putative glycerol-3-phosphate dehydrogenase (gpdh) | #N/A | 0 | 0 | 0 | 3 |
| TGME49_304870 | #N/A | #N/A | 0 | 0 | 0 | 3 |
| TGME49_299110 | cleft lip and palate transmembrane protein 1 (clptm1) pr | #N/A | 0 | 0 | 0 | 3 |
| TGME49_286740 | microneme-like protein | #N/A | 0 | 0 | 3 | 0 |
| TGME49_279350 | hypothetical protein | #N/A | 0 | 0 | 3 | 0 |
| TGME49_278510 | protein phosphatase 2C domain-containing protein | #N/A | 0 | 0 | 0 | 3 |
| TGME49_276130 | cathepsin CPC2 | TgCPC2 | 0 | 0 | 0 | 3 |
| TGME49_267400 | ribosomal protein RPL32 | #N/A | 0 | 0 | 0 | 3 |
| TGME49_262430 | 4-hydroxy-3-methylbut-2-en-1-yl diphosphate synthase | #N/A | 0 | 0 | 0 | 3 |
| TGME49_259260 | membrane protein FtsH1 | #N/A | 0 | 0 | 0 | 3 |
| TGME49_257530 | transporter, major facilitator family protein | #N/A | 0 | 0 | 0 | 3 |
| TGME49_257380 | hypothetical protein | #N/A | 0 | 0 | 3 | 0 |
| TGME49_251770 | hypothetical protein | #N/A | 0 | 0 | 3 | 0 |
| TGME49_231640 | alveolin domain containing intermediate filament IMC1 | #N/A | 0 | 0 | 0 | 3 |
| TGME49_226860 | SAG-related sequence SRS67 | #N/A | 0 | 0 | 0 | 3 |
| TGME49_226420 | peptidase family M3 protein | #N/A | 0 | 0 | 0 | 3 |
| TGME49_225120 | hypothetical protein | #N/A | 0 | 0 | 0 | 3 |
| TGME49_221310 | aminopeptidase N protein | #N/A | 0 | 0 | 0 | 3 |
| TGME49_220950 | hypothetical protein | MAF1 | 0 | 0 | 3 | 0 |
| TGME49_216630 | putative trigger factor protein | #N/A | 0 | 0 | 0 | 3 |
| TGME49_213010 | hypothetical protein | #N/A | 0 | 0 | 0 | 3 |
| TGME49_202010 | hypothetical protein | #N/A | 0 | 0 | 3 | 0 |
| TGME49_321620 | dynamin-related protein DRPB | #N/A | 0 | 0 | 0 | 2 |
| TGME49_318460 | P-type ATPase of unknown pump specificity (type V) prot | #N/A | 0 | 0 | 0 | 2 |
| TGME49_318440 | helicase associated domain (ha2) protein | #N/A | 0 | 0 | 0 | 2 |
| TGME49_314890 | ThiF family protein | #N/A | 0 | 0 | 0 | 2 |
| TGME49_313330 | rhoptry kinase family protein ROP27 | #N/A | 0 | 0 | 0 | 2 |
| TGME49_305980 | pyruvate dehydrogenase complex subunit PDH-E3I | #N/A | 0 | 0 | 0 | 2 |
| TGME49_301450 | FG-GAP repeat-containing protein | #N/A | 0 | 0 | 2 | 0 |
| TGME49_297500 | T-complex protein 1 eta subunit | #N/A | 0 | 0 | 0 | 2 |
| TGME49_296010 | phosphatidylinositol 3- and 4-kinase | #N/A | 0 | 0 | 2 | 0 |
| TGME49_293430 | hypothetical protein | #N/A | 0 | 0 | 2 | 0 |
| TGME49_291140 | CCR4-Not complex component, Not1 protein | #N/A | 0 | 0 | 0 | 2 |
| TGME49_290678 | hypothetical protein | #N/A | 0 | 0 | 0 | 2 |
| TGME49_285830 | hypothetical protein | #N/A | 0 | 0 | 0 | 2 |
| TGME49_281675 | putative protein kinase | ROP45 | 0 | 0 | 2 | 0 |
| TGME49_276210 | phosphoglycerate mutase family protein | #N/A | 0 | 0 | 0 | 2 |
| TGME49_272390 | hypothetical protein | #N/A | 0 | 0 | 0 | 2 |
| TGME49_272380 | hypothetical protein | #N/A | 0 | 0 | 0 | 2 |
| TGME49_269450 | hypothetical protein | #N/A | 0 | 0 | 0 | 2 |
| TGME49_268880 | hypothetical protein | #N/A | 0 | 0 | 2 | 0 |
| TGME49_268330 | hypothetical protein | #N/A | 0 | 0 | 0 | 2 |
| TGME49_264720 | hypothetical protein | #N/A | 0 | 0 | 2 | 0 |
| TGME49_264440 | signal recognition particle receptor beta subunit protein | #N/A | 0 | 0 | 0 | 2 |
| TGME49_257170 | hypothetical protein | #N/A | 0 | 0 | 0 | 2 |
| TGME49_255660 | EF hand domain-containing protein | #N/A | 0 | 0 | 0 | 2 |
| TGME49_253170 | putative zinc carboxypeptidase | #N/A | 0 | 0 | 0 | 2 |
| TGME49_249560 | DNA-directed RNA polymerase alpha chain rpoA | #N/A | 0 | 0 | 0 | 2 |

| | | | | | | |
|---|---|---|---|---|---|---|
| TGME49_244250 | hypothetical protein | #N/A | 0 | 0 | 2 | 0 |
| TGME49_236890 | hypothetical protein | GRA37 | 0 | 0 | 2 | 0 |
| TGME49_234280 | AMP deaminase | #N/A | 0 | 0 | 2 | 0 |
| TGME49_232180 | UBA/TS-N domain-containing protein | #N/A | 0 | 0 | 0 | 2 |
| TGME49_226660 | hypothetical protein | #N/A | 0 | 0 | 2 | 0 |
| TGME49_219540 | cytosolic tRNA-Ala synthetase | #N/A | 0 | 0 | 0 | 2 |
| TGME49_214620 | putative hypoxia- inducible factor prolyl hydroxylase (phd | #N/A | 0 | 0 | 2 | 0 |
| TGME49_213520 | peptidase M20D, amidohydrolase | #N/A | 0 | 0 | 0 | 2 |
| TGME49_203990 | rhoptry protein ROP12 | #N/A | 0 | 0 | 0 | 2 |
| TGME49_202200 | hypothetical protein | #N/A | 0 | 0 | 2 | 0 |
| TGME49_278830 | glucose-6-phosphate 1-dehydrogenase | #N/A | 0 | 2 | 72 | 77 |
| TGME49_269920 | phosphatidylserine decarboxylase | PSD1 | 0 | 3 | 42 | 55 |
| TGME49_311470 | rhoptry neck protein RON5 | #N/A | 0 | 6 | 90 | 97 |
| TGME49_252360 | rhoptry kinase family protein ROP24 (incomplete catalyti | #N/A | 0 | 2 | 29 | 23 |
| TGME49_269190 | glyceraldehyde-3-phosphate dehydrogenase GAPDH2 | #N/A | 6 | 4 | 107 | 128 |
| TGME49_277240 | #N/A | NTPase I | 0 | 2 | 18 | 24 |
| TGME49_244560 | putative heat shock protein 90 | #N/A | 18 | 8 | 211 | 277 |
| TGME49_311720 | chaperonin protein BiP | #N/A | 22 | 19 | 355 | 362 |
| TGME49_310750 | emp24/gp25L/p24 family protein | #N/A | 3 | 3 | 41 | 51 |
| TGME49_215785 | #N/A | #N/A | 0 | 4 | 22 | 27 |
| TGME49_262050 | rhoptry kinase family protein ROP39 | ROP39 | 3 | 0 | 14 | 21 |
| TGME49_295125 | rhoptry protein ROP4 | #N/A | 0 | 4 | 26 | 20 |
| TGME49_294200 | glucose-6-phosphate 1-dehydrogenase | #N/A | 8 | 7 | 80 | 73 |
| TGME49_295110 | rhoptry protein ROP7 | #N/A | 4 | 9 | 45 | 42 |
| TGME49_223920 | rhoptry neck protein RON3 | #N/A | 17 | 14 | 72 | 94 |
| TGME49_308090 | rhoptry protein ROP5 | #N/A | 2 | 2 | 6 | 12 |
| TGME49_258580 | rhoptry protein ROP17 | #N/A | 2 | 4 | 12 | 14 |
| TGME49_275860 | hypothetical protein | GRA12B | 5 | 9 | 26 | 31 |
| TGME49_266760 | isocitrate dehydrogenase | #N/A | 2 | 2 | 6 | 8 |
| TGME49_219320 | acid phosphatase GAP50 | #N/A | 6 | 7 | 24 | 20 |
| TGME49_237250 | hypothetical protein | #N/A | 3 | 0 | 5 | 5 |
| TGME49_257680 | myosin light chain MLC1 | #N/A | 3 | 3 | 9 | 10 |
| TGME49_215775 | rhoptry protein ROP8 | #N/A | 3 | 5 | 8 | 14 |
| TGME49_276970 | hypothetical protein | #N/A | 3 | 0 | 3 | 3 |
| TGME49_220860 | DEAD/DEAH box helicase | #N/A | 2 | 0 | 0 | 4 |
| TGME49_289750 | ribosomal-ubiquitin protein RPL40 | #N/A | 9 | 4 | 14 | 11 |
| TGME49_219820 | putative polyubiquitin UbC | #N/A | 9 | 4 | 14 | 11 |
| TGME49_244320 | ribosomal protein RPL24 | #N/A | 3 | 2 | 4 | 5 |
| TGME49_310490 | ribosomal protein RPL27A | #N/A | 3 | 3 | 3 | 6 |
| TGME49_300000 | ribosomal protein RPL18 | #N/A | 13 | 12 | 9 | 25 |
| TGME49_229670 | ribosomal protein RPS23 | #N/A | 5 | 9 | 8 | 11 |
| TGME49_249390 | glutamate/leucine/phenylalanine/valine dehydrogenase | #N/A | 6 | 9 | 9 | 11 |
| TGME49_260240 | ccr4-associated factor family protein | #N/A | 3 | 0 | 0 | 4 |
| TGME49_273760 | heat shock protein HSP70 | #N/A | 47 | 41 | 52 | 65 |
| TGME49_320020 | transporter, major facilitator family protein | #N/A | 4 | 4 | 5 | 5 |
| TGME49_255190 | myosin C | #N/A | 4 | 4 | 6 | 4 |
| TGME49_227360 | ribosomal protein RPL3 | #N/A | 19 | 22 | 17 | 34 |
| TGME49_316400 | #N/A | #N/A | 10 | 12 | 12 | 15 |
| TGME49_288380 | heat shock protein HSP90 | #N/A | 23 | 19 | 20 | 31 |
| TGME49_263180 | myosin D | #N/A | 9 | 10 | 13 | 10 |
| TGME49_221620 | putative beta-tubulin | #N/A | 9 | 15 | 10 | 18 |
| TGME49_209600 | hypothetical protein | #N/A | 3 | 5 | 4 | 5 |
| TGME49_288720 | ribosomal protein RPL10 | #N/A | 7 | 22 | 12 | 18 |
| TGME49_235470 | myosin A | #N/A | 34 | 39 | 35 | 39 |
| TGME49_263700 | ribosomal protein RPS14 | #N/A | 19 | 16 | 11 | 24 |
| TGME49_263050 | ribosomal protein RPL13 | #N/A | 4 | 7 | 4 | 7 |
| TGME49_201680 | putative eukaryotic initiation factor-3 subunit 10 | #N/A | 2 | 3 | 2 | 3 |
| TGME49_290200 | NAD/NADP octopine/nopaline dehydrogenase, alpha-heli | #N/A | 2 | 2 | 0 | 4 |
| TGME49_227030 | hypothetical protein | #N/A | 3 | 0 | 0 | 3 |
| TGME49_228470 | ribosomal protein RPL15 | #N/A | 16 | 20 | 19 | 16 |
| TGME49_289530 | ribosomal protein RPL19 | #N/A | 10 | 8 | 10 | 7 |
| TGME49_253430 | putative asparagine synthetase | #N/A | 9 | 7 | 7 | 8 |
| TGME49_215460 | ribosomal protein RPS24 | #N/A | 8 | 8 | 4 | 11 |
| TGME49_204020 | ribosomal protein RPL8 | #N/A | 10 | 17 | 10 | 15 |
| TGME49_294800 | #N/A | #N/A | 42 | 49 | 35 | 48 |
| TGME49_286420 | #N/A | #N/A | 42 | 49 | 35 | 48 |
| TGME49_210690 | ribosomal protein RPS6 | #N/A | 5 | 10 | 7 | 6 |
| TGME49_284190 | pyruvate carboxylase | #N/A | 452 | 618 | 446 | 480 |
| TGME49_207440 | ribosomal protein RPS4 | #N/A | 19 | 24 | 19 | 18 |
| TGME49_248340 | GTP-binding nuclear protein ran/tc4 | #N/A | 17 | 16 | 11 | 17 |
| TGME49_289690 | glyceraldehyde-3-phosphate dehydrogenase GAPDH1 | #N/A | 26 | 38 | 21 | 33 |
| TGME49_236650 | DEAD (Asp-Glu-Ala-Asp) box polypeptide 17 | #N/A | 11 | 10 | 7 | 10 |
| TGME49_274060 | 2-oxoglutarate/malate translocase OMT | #N/A | 2 | 3 | 0 | 4 |

| | | | | | | |
|---|---|---|---|---|---|---|
| TGME49_220400 | actin depolymerizing factor ADF | #N/A | 3 | 2 | 2 | 2 |
| TGME49_248390 | ribosomal protein RPL26 | #N/A | 7 | 7 | 5 | 6 |
| TGME49_221320 | acetyl-CoA carboxylase ACC1 | #N/A | 1416 | 2120 | 1376 | 1400 |
| TGME49_218560 | acetyl-coA carboxylase ACC2 | #N/A | 488 | 626 | 404 | 463 |
| TGME49_270510 | asparaginyl-tRNA synthetase (NOB+tRNA synthase) | #N/A | 15 | 12 | 8 | 13 |
| TGME49_226250 | DEAD (Asp-Glu-Ala-Asp) box polypeptide DDX3X | #N/A | 4 | 5 | 2 | 5 |
| TGME49_292130 | ribosomal protein RPL13A | #N/A | 14 | 17 | 12 | 12 |
| TGME49_266960 | beta-tubulin | #N/A | 11 | 15 | 7 | 13 |
| TGME49_232300 | ribosomal protein RPS3 | #N/A | 22 | 19 | 15 | 16 |
| TGME49_247510 | fructose-bisphospatase II | #N/A | 15 | 9 | 7 | 11 |
| TGME49_262620 | RNA recognition motif-containing protein | #N/A | 11 | 9 | 8 | 7 |
| TGME49_305520 | ribosomal protein RPS2 | #N/A | 15 | 12 | 11 | 9 |
| TGME49_309590 | rhoptry protein ROP1 | #N/A | 6 | 9 | 5 | 6 |
| TGME49_223050 | ribosomal protein RPS20 | #N/A | 8 | 6 | 3 | 7 |
| TGME49_213350 | ribosomal protein RPS15 | #N/A | 5 | 5 | 3 | 4 |
| TGME49_205470 | putative translation elongation factor 2 family protein | #N/A | 7 | 3 | 0 | 7 |
| TGME49_245680 | ribosomal protein RPL21 | #N/A | 12 | 10 | 5 | 10 |
| TGME49_209030 | actin ACT1 | #N/A | 54 | 56 | 32 | 42 |
| TGME49_253290 | valyl-tRNA synthetase | #N/A | 3 | 3 | 0 | 4 |
| TGME49_289760 | hypothetical protein | #N/A | 31 | 28 | 13 | 26 |
| TGME49_306030 | glutathione s-transferase, n-terminal domain containing | #N/A | 5 | 6 | 2 | 5 |
| TGME49_312090 | ribosomal protein RPL23 | #N/A | 8 | 11 | 5 | 7 |
| TGME49_261570 | ribosomal protein RPL7A | #N/A | 5 | 11 | 4 | 6 |
| TGME49_204400 | putative ATPase synthase subunit alpha | #N/A | 10 | 6 | 6 | 4 |
| TGME49_312050 | putative small GTPase Rab2 | #N/A | 10 | 13 | 6 | 8 |
| TGME49_263040 | ribosomal protein RPS16 | #N/A | 3 | 2 | 0 | 3 |
| TGME49_250770 | putative eukaryotic initiation factor-4A | #N/A | 26 | 21 | 13 | 15 |
| TGME49_309120 | ribosomal protein RPL4 | #N/A | 16 | 20 | 7 | 14 |
| TGME49_320050 | ribosomal protein RPL5 | #N/A | 23 | 24 | 14 | 13 |
| TGME49_278870 | myosin F | #N/A | 22 | 31 | 18 | 12 |
| TGME49_245460 | ribosomal protein RPS8 | #N/A | 17 | 18 | 12 | 7 |
| TGME49_249900 | putative adenine nucleotide translocator | #N/A | 28 | 28 | 12 | 18 |
| TGME49_239100 | ribosomal protein RPS7 | #N/A | 10 | 15 | 5 | 8 |
| TGME49_249250 | ribosomal protein RPL35A | #N/A | 15 | 17 | 5 | 11 |
| TGME49_290670 | leucyl aminopeptidase LAP | #N/A | 12 | 6 | 2 | 7 |
| TGME49_239820 | D-3-phosphoglycerate dehydrogenase | #N/A | 7 | 9 | 2 | 6 |
| TGME49_248880 | GTPase RAB7 | #N/A | 6 | 6 | 2 | 4 |
| TGME49_289800 | hypothetical protein | #N/A | 0 | 4 | 0 | 2 |
| TGME49_226970 | ribosomal protein RPS11 | #N/A | 16 | 17 | 7 | 9 |
| TGME49_324600 | heat shock protein | #N/A | 11 | 14 | 5 | 7 |
| TGME49_232940 | #N/A | #N/A | 11 | 14 | 5 | 7 |
| TGME49_232710 | ribosomal protein RPS3A | #N/A | 9 | 8 | 2 | 6 |
| TGME49_284560 | ribosomal protein RPL9 | #N/A | 8 | 7 | 3 | 3 |
| TGME49_305510 | hypothetical protein | #N/A | 2 | 3 | 2 | 0 |
| TGME49_273460 | eukaryotic translation initiation factor 3 subunit 6 interac | #N/A | 8 | 8 | 0 | 6 |
| TGME49_240860 | acyltransferase domain-containing protein | #N/A | 5 | 3 | 0 | 3 |
| TGME49_313010 | DEAD (Asp-Glu-Ala-Asp) box polypeptide DDX6 | #N/A | 17 | 12 | 0 | 10 |
| TGME49_251780 | heat shock protein | #N/A | 5 | 7 | 0 | 4 |
| TGME49_242330 | ribosomal protein RPS5 | #N/A | 4 | 2 | 0 | 2 |
| TGME49_262380 | putative elongation factor Tu | #N/A | 4 | 6 | 0 | 3 |
| TGME49_230940 | hypothetical protein | #N/A | 9 | 2 | 0 | 3 |
| TGME49_262670 | ribosomal protein RPL18A | #N/A | 13 | 11 | 0 | 6 |
| TGME49_299050 | ribosomal protein RPL17 | #N/A | 4 | 4 | 0 | 2 |
| TGME49_309820 | ribosomal protein RPL11 | #N/A | 4 | 5 | 0 | 2 |
| TGME49_309752 | putative succinate-Coenzyme A ligase, beta subunit | #N/A | 5 | 4 | 0 | 2 |
| TGME49_288830 | NADH dehydrogenase (NDH2-II) | #N/A | 9 | 13 | 2 | 2 |
| TGME49_263300 | eukaryotic porin protein | #N/A | 9 | 8 | 0 | 3 |
| TGME49_314970 | root hair defective 3 gtp-binding protein (rhd3) protein | #N/A | 2 | 0 | 0 | 0 |
| TGME49_314810 | ribosomal protein RPL7 | #N/A | 2 | 3 | 0 | 0 |
| TGME49_314400 | putative pyruvate dehydrogenase E1 component, beta su | #N/A | 2 | 0 | 0 | 0 |
| TGME49_313670 | adaptin n terminal region domain-containing protein | #N/A | 0 | 2 | 0 | 0 |
| TGME49_311430 | fibrillarin | #N/A | 5 | 0 | 0 | 0 |
| TGME49_311290 | protein tyrosine phosphatase family protein, ptpla protei | #N/A | 2 | 0 | 0 | 0 |
| TGME49_301440 | calcium-dependent protein kinase CDPK1 | #N/A | 2 | 2 | 0 | 0 |
| TGME49_300140 | putative elongation factor 1-gamma | #N/A | 3 | 0 | 0 | 0 |
| TGME49_292080 | leucyl-tRNA synthetase | #N/A | 0 | 2 | 0 | 0 |
| TGME49_289300 | methionyl-tRNA synthetase | #N/A | 2 | 0 | 0 | 0 |
| TGME49_288360 | tryptophanyl-tRNA synthetase (TrpRS2) | #N/A | 2 | 0 | 0 | 0 |
| TGME49_287500 | putative T complex chaperonin | #N/A | 5 | 6 | 0 | 0 |
| TGME49_284050 | DEAD/DEAH box helicase domain-containing protein | #N/A | 4 | 6 | 0 | 0 |
| TGME49_270270 | hypothetical protein | #N/A | 2 | 0 | 0 | 0 |
| TGME49_269600 | biotin-requiring enzyme domain-containing protein | #N/A | 7 | 2 | 0 | 0 |
| TGME49_264080 | acyl carrier protein ACP | #N/A | 3 | 5 | 0 | 0 |

| | | | | | | |
|---|---|---|---|---|---|---|
| TGME49_262960 | putative U1 snRNP-associated protein Usp106 | #N/A | 2 | 2 | 0 | 0 |
| TGME49_248480 | ribosomal protein RPS9 | #N/A | 4 | 0 | 0 | 0 |
| TGME49_243950 | putative prohibitin | #N/A | 2 | 3 | 0 | 0 |
| TGME49_243570 | ribosomal protein RPS26 | #N/A | 0 | 3 | 0 | 0 |
| TGME49_239760 | ribosomal protein RPL22 | #N/A | 2 | 0 | 0 | 0 |
| TGME49_236570 | lysine decarboxylase family protein | #N/A | 2 | 2 | 0 | 0 |
| TGME49_235930 | domain K- type RNA binding proteins family protein | #N/A | 2 | 0 | 0 | 0 |
| TGME49_232230 | ribosomal protein RPL30 | #N/A | 4 | 2 | 0 | 0 |
| TGME49_229360 | transaldolase | #N/A | 2 | 0 | 0 | 0 |
| TGME49_229250 | #N/A | #N/A | 0 | 2 | 0 | 0 |
| TGME49_227600 | ribosomal protein RPL34 | #N/A | 2 | 5 | 0 | 0 |
| TGME49_222900 | phosphoserine phosphatase | #N/A | 0 | 2 | 0 | 0 |
| TGME49_221380 | alba 1 | #N/A | 3 | 5 | 0 | 0 |
| TGME49_219720 | putative Ras-related protein Rab-5C | #N/A | 5 | 0 | 0 | 0 |
| TGME49_218280 | putative eukaryotic porin | #N/A | 2 | 2 | 0 | 0 |
| TGME49_217570 | ribosomal protein RPS27 | #N/A | 0 | 2 | 0 | 0 |
| TGME49_216880 | guanine nucleotide-binding protein | #N/A | 4 | 0 | 0 | 0 |
| TGME49_216860 | DEAD (Asp-Glu-Ala-Asp) box polypeptide DDX39 | #N/A | 4 | 0 | 0 | 0 |
| TGME49_214770 | putative small GTP binding protein rab1a | #N/A | 0 | 3 | 0 | 0 |
| TGME49_213910 | hypothetical protein | #N/A | 0 | 2 | 0 | 0 |
